# A Resource for Assessing Information Processing in the Developing Brain Using EEG and Eye Tracking

**DOI:** 10.1101/092213

**Authors:** Nicolas Langer, Erica J. Ho, Lindsay M. Alexander, Helen Y. Xu, Renee K. Jozanovic, Simon Henin, Samantha Cohen, Enitan T. Marcelle, Lucas C. Parra, Michael P. Milham, Simon P. Kelly

**Author notes:** These authors contributed equally to this work (Nicolas Langer, Erica J. Ho). Corresponding author(s): Nicolas Langer, Michael P. Milham and Simon P. Kelly.

## Abstract

We present a dataset combining electrophysiology and eye tracking intended as a resource for the investigation of information processing in the developing brain. The dataset includes high-density task-based and task-free EEG, eye tracking, and cognitive and behavioral data collected from 126 individuals (ages: 6–44). The task battery spans both the simple/complex and passive/active dimensions to cover a range of approaches prevalent in modern cognitive neuroscience. The active task paradigms facilitate principled deconstruction of core components of task performance in the developing brain, whereas the passive paradigms permit the examination of intrinsic functional network activity during varying amounts of external stimulation. Alongside these neurophysiological data, we include an abbreviated cognitive test battery and questionnaire-based measures of psychiatric functioning. We hope that this dataset will lead to the development of novel assays of neural processes fundamental to information processing, which can be used to index healthy brain development as well as detect pathologic processes.

## BACKGROUND AND SUMMARY

It has become increasingly apparent that there are abundant links between cognitive deficits and mental health disorders. Progress towards identifying these relationships has been limited on the clinical side by validity and specificity issues pervading conventional diagnostic categories, and on the neuroscientific side by the restricted scope of conventional datasets (Kapur et al., 2012; Kozak and Cuthbert, 2016). These realities have spurred an imperative for clinically-focused cognitive neuroscience research to move from the confined study of specific facets of cognition in specific diagnostic groups, towards the ideal of examining all facets in all individuals (Insel and Cuthbert, 2015). The National Institute of Mental Health has taken a leading role in these efforts by establishing the Research Domain Criteria (RDoC) Project as a framework for forging multidimensional characterizations of mental illness. A central aspect of this framework is the integration of information across multiple levels (e.g. from genetics to self-report), and the recognition of human neuroimaging and neurophysiology measures as potentially providing key dimensions (Cuthbert and Insel, 2013).

The field of cognitive neuroscience encompasses a wide breadth of approaches to measuring functionally relevant neural activity, each with its own pros and cons. For example, some research has focused on neural activity measurements during task performance, which can link discrete neural signatures to behavioral outcomes recorded simultaneously. Meanwhile, “task-free” neural recordings taken during resting or passive stimulation conditions (e.g., naturalistic viewing) have increased in popularity because they provide a broader view on neural dynamics that transcend specific, circumscribed task scenarios. Additionally, they remove behavioral requirements that can at times limit their utility in developing and clinical populations. Another principal dimension along which approaches vary greatly is that of paradigm complexity. This dimension involves a definite trade-off: on one hand, elementary tasks involving reduced stimuli with few attributes and simple action mappings afford greater possibilities to link low-resolution neural activity measures to well-defined computations, partly because the computational building blocks of a simple task are more easily identified. On the other hand, such reduced, simplified tasks correspond to artificial behavioral scenarios, and it is important to measure neural activity during more complex, ecologically valid behavioral scenarios that lie closer both to real-life behavior and to clinical symptomology.

Here, we present a novel battery of EEG-based paradigms that attempts to “run the gamut” in both of these respects, widely spanning both the passive and active, as well as simple and complex paradigm dimensions that typically distinguish approaches at the far corners of cognitive neuroscience. This approach embraces the concept of integrating across levels. The battery includes three active task paradigms, which allow, to varying degrees, principled deconstruction of core components of task performance (see Table 1). The simplest paradigm permits the tracing of the three major processing stages for simple contrast decisions (O’Connell et al., 2012). The second paradigm involves the learning of simple sequences (Steinemann et al., 2016). The third emulates a standard neuropsychological processing speed task, which involves multiple perceptual decisions, short-term memory, and motor responses. For all of these tasks, simultaneous eye tracking provides a rich complement to EEG-based characterizations of neural processing and cognition.

**Table 1.** Experimental Paradigms Included. An overview of the six EEG and eye tracking paradigms.

| Task | Depth of Processing/Degree of Stimulation | Description |
| --- | --- | --- |
| **Active (Task-Dependent) Paradigms** |  |  |
| Contrast change | Minimal | Probes basic elements of sensorimotor translations, e.g. sensory evidence encoding, decision formation and motor preparation, providing dynamic measurements of each processing stage in isolation. |
| Sequence learning | Moderate | Assesses successive visuo-spatial sequence learning by using semantically unloaded stimuli, track the progress of gradual memory formation |
| Symbol search | Complex | A computerized version of a clinical pediatric assessment measuring processing speed capacity in a visual search task, which involves multiple perceptual decisions, short-term memory and motor response. |
| **Passive (Task-Independent) Paradigms** |  |  |
| Resting-state | None | Measures endogenous brain activity during rest. |
| Surround suppression | Minimal | Measures excitatory (using the steady-state visually evoked potential; SSVEP) and inhibitory (using the surround-suppression effect) neurophysiological activity during sensory processing with semantically unloaded stimuli. |
| Naturalistic viewing | Complex | Measures neurophysiological activity during higher-level audio-visual stimulation (movies). |

The battery also includes three passive paradigms, which permit the examination of intrinsic functional network activity during different amounts of external stimulation, namely, no stimulation (classical resting-state); simple and reduced (surround-suppression paradigm); and complex and rich (videos; Table 1). Whereas the simpler stimulation offers insight into elemental facets of information processing such as excitatory/inhibitory balance, the complex video stimuli allow measurement of engagement with naturalistic content (Dmochowski et al., 2014).

Alongside these neurophysiological data, abbreviated, standardized tests of intelligence and academic achievement, and self- or parent-reported measures of psychiatric functioning have been included.

In our initial release, we present high-density task-based and task-free EEG, eye tracking, and cognitive and behavioral data for 126 subjects ages 6–44, the majority of whom do not have a history of clinical illness. Our long-term goal is to collect data on this multi-level, multi-modal battery from a diverse community sample, including patient populations. Ultimately, we hope that this dataset will provide a rich new set of metrics for assaying neural processes fundamental to perception and cognition across a continuum from healthy to pathological functioning, and thereby contribute to understanding and better diagnosing a broad range of brain pathologies.

## METHODS

### Participants and experiment overview

126 Individuals between the ages of 6 and 44 were invited to participate in a study investigating domain-general cognitive processes related to attention, working memory, perception, and decision-making across a range of task/stimulation contexts. The participants were recruited from both the Child Mind Medical Practice, as well as the wider New York City-area community. 80.2% were typically developing, and 19.8% were diagnosed with one or more clinical disorders (see Table 2 for a summary of diagnostic categories represented in the sample). The participants were 54.8% male, 45.2% female; 45.2% identified as Black or African American, 32.7% as White,.04% as Asian, and 17.3% as other race or races (see Figure 1). Also included are measures of subject handedness (Annett, 1970) and socioeconomic status (MacArthur Scale of Subjective Social Status, http://www.macses.ucsf.edu/research/socialenviron/sociodemographic.php).

**Table 2.** Diagnosis Status. Diagnoses of subjects are shown, spanning 11 categories of diagnoses, in addition to no diagnosis. Frequency and percentages are shown. Note that subjects may have single or multiple diagnoses.

| Diagnostic Category | Frequency | % of clinical sample | % of total sample |
| --- | --- | --- | --- |
| No Diagnosis | 101 | N/A | 0.80 |
| Attention | 12 | 0.48 | 0.10 |
| Anxiety | 10 | 0.40 | 0.08 |
| Learning | 7 | 0.28 | 0.06 |
| OCD | 4 | 0.16 | 0.03 |
| ASD | 2 | 0.08 | 0.02 |
| Depressive | 2 | 0.08 | 0.02 |
| Trauma | 2 | 0.08 | 0.02 |
| Disruptive | 2 | 0.08 | 0.02 |
| Motor | 2 | 0.08 | 0.02 |
| Language | 1 | 0.04 | 0.01 |
| Mood | 1 | 0.04 | 0.01 |

**Figure 1.**
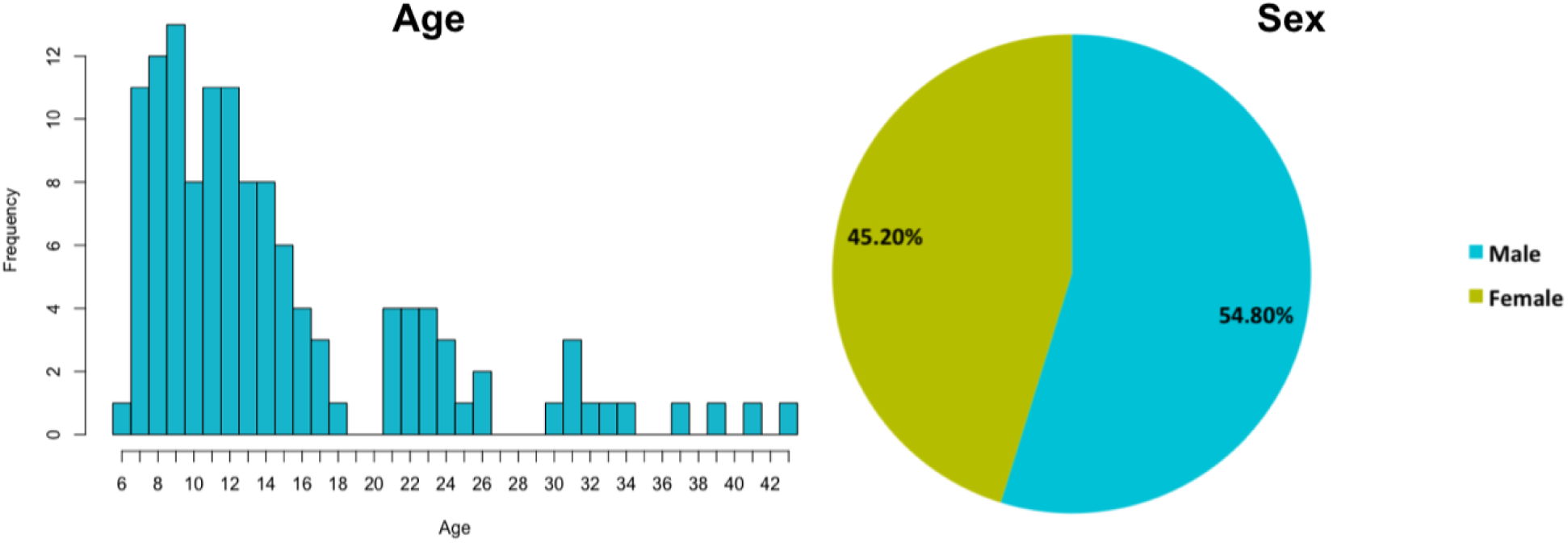
Age and Sex. Age distribution of subjects is displayed on the left plot. Ages ranged from 6 to 44. Sex breakdown of participants is displayed on the right plot.

Prior to visiting the laboratory, participants (or their legal guardians, in the case of participants under the age of 18) completed a 10 min. pre-screening interview over the phone with a research assistant to confirm their eligibility and safety to participate in the study. This brief interview obtained information regarding an individual’s psychiatric history, including past or present diagnoses and/or treatment, as well as current medications and any neurological disorders. If a participant demonstrated no contraindications for EEG (e.g., history of seizures or epilepsy), he or she was then scheduled for a research study appointment.

The full battery of EEG and eye tracking tasks and behavioral assessments was five hours in duration; participants were permitted to split their visit into two shorter sessions, lasting 3 hours (EEG recording and eye tracking portion) and 2 hours (cognitive and behavioral assessment portion) respectively. For those who elected to participate in the single, full-length session, the EEG and eye tracking tasks always preceded the cognitive and behavioral testing.

The study was approved by the Chesapeake Institutional Review Board. Written informed consent was obtained from all participants or their legal guardians prior to the start of the experiment; additionally, written assent was obtained from participants under the age of 18 and over the age of 6. Consent was also obtained for data sharing through the 1000 Functional Connectomes Project [http://fcon_1000.projects.nitrc.org/].

### Behavioral/Cognitive Assessments

All behavioral and cognitive assessments are described in Table 3.

#### Behavioral

Behavioral self-report measures were acquired via the online Self-Assessment Portal of the Collaborative Informatics and Neuroimaging Suite (COINS).

#### Cognitive

Cognitive testing was administered by trained research assistants in a sound-shielded room. The participants’ responses were first-scored by the research assistant who administered the test; then, to ensure accuracy, the entire set of responses were again scored by another trained research assistant. Furthermore, all test scores were double-entered into the database by two different research assistants. Both raw scores and standard scores are provided as part of this dataset.

### Data Acquisition Overview

Participants were seated in a sound-attenuated and dark experiment room at a distance of 70 cm from a 17-inch CRT monitor (resolution 800 x 600 pixels, vertical refresh rate of 100 Hz). A stable head position was ensured via the chin rest. Subjects were instructed to stay as still as possible during the tasks. Two breaks were included in the EEG session, during which electrode impedance levels were checked and reduced if necessary. Participants were also offered snacks and juice during the breaks, and encouraged to rest.

Stimulus presentation was programmed in MATLAB (6.1, The Math-Works, Natick, MA, 2000), using the PsychToolbox extension (Brainard, 1997; Pelli, 1997). The order of the EEG and eye tracking paradigms was the same for all participants. Instructions for the tasks were presented on the computer screen, and a research assistant answered questions from the participant from the adjacent control room through an intercom. Compliance with the task instructions was confirmed through a live video-feed to the control room. If participants were approximately 12 years of age or younger, they were joined in the experiment room by an additional research assistant who proctored their testing session; otherwise, participants completed the EEG and eye tracking tasks siting alone in the room.

#### EEG Acquisition

High-density EEG data were recorded at a sampling rate of 500 Hz with a bandpass of 0.1 to 100 Hz, using a 128-channel EEG Geodesic Hydrocel system. The recording reference was at Cz (vertex of the head). For each participant, head circumference was measured and an appropriately sized EEG net was selected. The impedance of each electrode was checked prior to recording, to ensure good contact, and was kept below 40 kOhm. Time to prepare the EEG net was no more than 30 minutes. Impedance was tested every 30 minutes of recording and saline added if needed.

#### Eye Tracking Acquisition

During all the EEG paradigms, eye position and pupil dilation were recorded binocularly with an infrared video-based eye tracker (iView-X Red-m, SMI GmbH; http://www.smivision.com/en.html) at a sampling rate of 120 Hz and an instrument spatial resolution of a nominal 0.1°. The eye tracker was calibrated with a 5-point grid before each paradigm.

### Paradigm Overview

Both task-independent (passive) and task-based (active) paradigms were included in the EEG battery as they play complementary roles in the comprehensive investigation of human brain function. Paradigms were also selected to vary widely in the degree of sensory stimulation involved and/or the depth of processing, from simple to complex. Our task-free paradigms permit examination of intrinsic functional network responses to different degrees of external stimulation, e.g., no stimulation (classical resting-state); simple and reduced (surround-suppression paradigm); and complex and rich (videos). In general, such passive paradigms enable measurement of neurophysiological indices of brain function on a relatively equal footing across a wider population, including low-functioning neurological and psychiatric populations for whom task-based assays are a challenge. On the other hand, our task-based paradigms aim to isolate distinct, fundamental information processing steps that play a core role in most neuropsychological and psychometric assessments, and thereby furnish a systems-level, neurophysiologically-based account of the factors underlying observed impairments in accuracy and/or response speed. Taken together, our EEG paradigm battery is intended to provide a window into neurophysiological mechanisms underlying domain-general cognitive functions, which account for a diverse range of behaviors and should thus, in theory, be possible to connect with psychiatric symptoms.

Note that the purpose of the “Output Measures” section in each of the following paradigm descriptions is to propose or guide possible analysis strategies for other researchers, without any intention to restrict the scope for using the present dataset in further creative and distinctive ways.

#### Paradigm #1 (Passive): Resting-State

##### Task Overview

The acquisition of endogenous brain activity without any external stimulation has become very popular in the EEG and functional MRI communities. The low cognitive demand and relatively short duration of resting-state recordings make them well suited for studying pediatric and clinical populations with low tolerance for standard paradigms and acquisitions (Fox and Greicius, 2010). A growing number of studies have shown that many of the brain areas engaged during various cognitive tasks also form coherent large-scale brain networks that can be readily identified in data recorded during rest (Biswal et al., 1995; Damoiseaux et al., 2006; Smith et al., 2009).

Numerous studies have demonstrated high intra-individual stability for resting EEG measures (Vogel, 2000; Posthuma et al., 2001; Orekhova et al., 2003; Ivonin et al., 2004; Linkenkaer-Hansen et al., 2007). For example, it was demonstrated that individual participants could be identified based only on their resting EEG measures with a sensitivity as high as 88% and specificity of 99.5% (Napflin et al., 2007). Intraclass correlation coefficients have been used to show strong retest reliabilities for power in the alpha (8-14 Hz) and beta (15–30 Hz) bands, which ranged from r = 0.8 to r > 0.9 (Kondacs and Szabo, 1999). Finally, (Deuker et al., 2009) demonstrated the reproducibility of graph metrics of human brain functional networks obtained by resting-state EEG data. Collectively, these results suggest that resting-state EEG is highly reliable and thus can potentially provide stable biological markers that can be related to cognitive performance across individuals.

##### Stimuli & Experimental Design

Participants viewed a standard fixation cross in the center of the computer screen. The recorded voice of a female research assistant instructed them to “now open your eyes” (rest with eyes open for 20 s) and “now close your eyes” (rest with eyes closed for 40 s); this procedure was repeated 5 times, alternating between eyes opened and eyes closed. For purposes of analysis, we were mainly interested in the eyes-closed condition, due to the lower frequency of eye blink and muscular artifacts. However, we interspersed the brief eyes-open blocks throughout the task in order to ensure that participants remained engaged for the duration of the task session.

##### Participant Instructions

“Fixate on the central cross. Open or close your eyes when you hear the request for it. Press to begin.”

##### Output Measures

There are various ways to analyze resting-state EEG data. One can examine the data in the frequency domain using classical power spectral analysis, which has been successfully employed to characterize subjects’ age (John et al., 1980), state of arousal (Borbely and Achermann, 1999), the presence of neurological or psychiatric disorders (John et al., 1988; Jeste et al., 2015; Kitsune et al., 2015; Loo et al., 2015), or task demands (Fernandez et al., 1995; Gevins et al., 1997). Advanced research on resting-state EEG and fMRI offers a novel approach for understanding synchronization of intrinsic fluctuations in neurophysiological activity, which is measured as a dependency between time-series obtained from different regions in the brain (Friston et al., 1993; Pascual-Marqui, 2007; De Vico Fallani et al., 2010; Langer et al., 2012). This includes frequency-domain analyses such as the characterization of global and local connectivity between EEG sources (i.e., functional- and effective-connectivity; graph theoretical network properties). Several researchers have also emphasized the value of investigating resting-state data from a temporal-spatial perspective to reveal microstates, which are stable spatial configurations of the electric field that vary across time (Lehmann et al., 1987; Pascual-Marqui et al., 1995; Murray et al., 2008). These spatially stationary microstates have been proposed to reflect basic building blocks of information processing (Lehmann et al., 1998).

#### Paradigm #2 (Passive): Surround Suppression

##### Task Overview

The surround suppression paradigm enables measurement of basic sensory excitation by visual stimuli and the suppressive contextual influence of the visual background, thereby providing insight into relative levels of excitability and inhibition in the human cortex. In this paradigm, periodic, visual, on-off flicker stimulation is used to elicit periodic EEG/MEG responses at the exact frequency of stimulation and its harmonics, known as the steady state visual evoked potential (SSVEP) (Regan, 1966; Regan, 1989). Being spectrally restricted to a single frequency, SSVEPs provide a measure of visual neural response amplitude with a higher signal-to-noise ratio than standard transient evoked potential approaches (Regan, 1966; Regan, 1989). SSVEP amplitude and phase can be measured to probe sensory sensitivity and latency (timing) information, respectively, and these measures can further be tracked over time to gain insight into dynamic aspects of sensory responses such as adaptation and attention orienting (Vanegas et al., 2015). SSVEP amplitude and topographic variation across individuals correlate with intelligence (Van Rooy et al., 2001) and depend on age (Macpherson et al., 2009). They have also been informative in the study of cognitive disorders such as schizophrenia, anxiety, stress, and epilepsy (Vialatte et al., 2010).

In our surround suppression paradigm, we present “foreground” flicker stimuli at a range of contrasts to probe basic visual excitation, and we also manipulate the contrast of a static surround pattern to probe basic inhibition. Surround suppression is the well-known phenomenon whereby the neural response to a delimited stimulus is suppressed by stimulation in the surrounding area, which has been widely observed in animal neurophysiology (e.g., Levitt and Lund, 1997; Cavanaugh et al., 2002), and in human psychophysics (e.g., Xing and Heeger, 2000), neuroimaging (e.g., Zenger-Landolt and Heeger, 2003), and electrophysiology (Vanegas et al., 2015). In our paradigm we obtain an index surround suppression by measuring the reduction in “foreground” SSVEP amplitude that results from the presence of the static surround. Surround suppression has become increasingly relevant in clinical research, with clear abnormalities reported in a range of disorders such as depression (Golomb et al., 2009), autism (Foss-Feig et al., 2013), schizophrenia (Dakin et al., 2005; Seymour et al., 2013), and migraine (Battista et al., 2011).

##### Stimuli & Experimental Design

We used the paradigm developed by Vanegas et al. (2015), adapted to include a restricted set of conditions that were established to provide the most robust measures. In each sequence of discrete 2.4 sec trials, four circular “foreground” stimuli (vertical grating, radius 2°) were flickered on-and-off at 25 Hz, embedded in a static (non-flickering) full-screen “surround” (see Figure 3). On the basis of recent work in which we demonstrated dramatic improvements in SSVEP signal-to-noise ratio (SNR), we flickered the upper discs with opposite temporal phase relative to the lower discs in the foreground, causing oscillatory summation on the scalp because of the cortical surface orientation of early retinotopic visual areas (Vanegas et al., 2013). Each trial began with the presentation of the fixation spot for 500 ms, after which the foreground and surround stimuli were simultaneously presented for 2400 ms. After an inter-trial interval of 500ms, the following trial was initiated. Foreground and surround patterns were sinusoidal luminance-modulated gratings with a spatial frequency of 1 cycle per degree in all conditions (see Fig 1A, B, and C in Vanegas et al., 2015). Across trials, we randomly varied foreground contrast (0%, 30%, 60% or 100%), surround contrast (0% or 100%) and surround orientation (parallel or orthogonal to the foreground, i.e., vertical or horizontal). Eye gaze was monitored continually using the eye tracker. The entire task was recorded in two blocks, each consisting of 64 trials and lasting ~3.6 mins.

##### Participant Instructions

“Just maintain fixation on the central spot at all times. Press to begin. First, we have to measure the position of your eyes. Just follow the circle with your eyes.”

##### Output Measures

The flickering foreground elicits a steady-state visual evoked potential (SSVEP) in the EEG over posterior scalp at the fundamental frequency of stimulation, the amplitude of which increases monotonically with foreground contrast (Lauritzen et al., 2010). Surround suppression is measured as a relative reduction in amplitude of the SSVEP due to surround contrast. As mentioned above, SSVEP amplitude and phase can also by tracked over time to examine temporal aspects of gain control as well as latency effects. These measures have the potential to provide a marker of improperly balanced excitation and inhibition in children with developmental disorders, as has been implicated in recent studies of autism (Foss-Feig et al., 2013).

#### Paradigm #3 (Passive): Naturalistic Stimuli

##### Task Overview

In recent years, there has been a significant expansion in the scope of studies utilizing naturalistic viewing paradigms (Bartels and Zeki, 2004; Hasson et al., 2004; Hasson et al., 2010). Naturalistic viewing paradigms, such as movies, have been shown to evoke patterns of neural activity that are synchronized across individuals, and even across species (Hasson et al., 2004; Hasson et al., 2008). In addition, time courses derived from features of the movie such as luminance and sound intensity can be used to investigate different facets of neurofunctional systems with improved precision. Movies thus provide a powerful and flexible medium through which to engage multiple networks in a concerted and dynamic fashion. From a clinical standpoint, the use of movies in the context of functional connectivity allows shorter data collection times and decreases head movement in both adults and children (Vanderwal et al., 2015).

The goal of the present paradigm was to measure variable engagement based on the strength of higher-level audio-visual responses, and to aid the understanding of the modulation of perception across ages and developmental stages (Petroni et al., 2016). Participants viewed 4 short, age-appropriate video clips taken from television and movies. There is evidence that children’s performance on reading, school readiness, and creativity tests improve after viewing educational programs such as *Sesame Street* (Cantlon and Li, 2013). Thus, the content of educational videos, such as those used in the current study, can interact with children’s school-based knowledge. These advantages of the natural viewing stimuli over a more traditional task with simple stimuli suggest that naturalistic studies of brain activity with real-world stimuli could serve as an important complement to highly controlled EEG paradigms.

##### Stimuli & Experimental Design

Participants viewed 4 short, age-appropriate video clips taken from television and movies. Each clip was between 2 and 6 min in length, for a total of 12:50 minutes.

(Prior to this task, parents were given the opportunity to review the full list of clips and exclude any video clips they deemed unsuitable for their children; no parents had any objections to the clips.). The following are a description of clips that we included in the *Naturalistic Stimuli Paradigm.*

<u>E-How video: How to Improve at Simple Arithmetic: Lessons in Math</u>

Rating: No parental guideline rating

Description: A female instructor introduces addition and multiplication tricks.

Rationale: This clip is included to probe for attention related difficulties.

Link: http://www.youtube.com/watch?v=pHoE7AMtXcA

Length: 1:40

<u>MIT K-12: “Fun with Fractals”</u>

Rating: No parental guideline rating

Description: This video depicts fractal-based geometry in everyday objects and visually depicts how some fractals are created.

Rationale: This clip is included to probe for attention related difficulties.

Link: http://www.youtube.com/watch?v=XwWyTts06tU

Length: 4:40

<u>Diary of a Wimpy Kid Trailer</u>

Rating: Rated PG for some rude humor and language

Description: This comedic movie trailer is a hyperbolic depiction of a child’s experience of middle school. It contains several character vignettes.

Rationale: This clip is included to probe for socially related anxiety.

Link: http://www.youtube.com/watch?v=7ZVEIgPeDCE

Length: 2:00

<u>Despicable Me</u>

Rating: Rated PG for rude humor and mild action

Description: In this animation, a new adoptive father reads his three children a bedtime story.

Rationale: This clip is included to probe for attachment formation related issues.

Link: http://www.youtube.com/watch?v=HNXxJIhVALI

Length: 2:50

##### Participant Instructions

“Now you can watch video clips. Enjoy! First, we have to measure the position of your eyes. Just follow with your eyes the circle. Press to begin.”

##### Output Measures

Naturalistic audiovisual stimuli have been shown to elicit highly reliable neural activity across multiple viewers (Hasson et al., 2004), with the level of such inter-subject correlation (ISC) linked to successful memory encoding (Hasson et al., 2008), and effective communication between individuals (Stephens et al., 2010). ISC usually is increased during scenes marked by high arousal and negative emotional valence (Hasson et al., 2004), and is strongest for familiar and naturalistic events (Hanson et al., 2009). Here, the EEG data were analyzed using Correlated Component Analysis (CCA) in order to parse relative inter-subject correlations (ISC). We are mainly interested in the similarity of neural response across subjects for naturalistic stimuli experienced in everyday life. To determine the neural similarity among subjects in response to a stimulus, the inter-subject correlation (ISC) of the EEG signal was calculated. The procedure is described in detail in previous studies (Dmochowski et al., 2012; Dmochowski et al., 2014).

In brief, the ISC is a measure of correlation among a group of subjects; larger values imply more similarity of the EEG signal across subjects in response to identical stimuli. The advantage of the ISC technique compared to averaging multiple trials is that it can be calculated with a single presentation of a novel stimulus, allowing naturalistic settings with continuous stimulation rather than discrete events (Zacks and Tversky, 2001; Fontanini and Katz, 2008; Ben-Yakov et al., 2012). The technique, based on the correlated component analysis, identifies linear combinations of electrodes—called components—that maximize the correlation across subjects. In general terms, CCA is very similar to a PCA, but rather than maximizing variance, it maximizes correlation between subjects (datasets). The technique has been described in detail in (Cohen and Parra, 2016) and applied on the data reported here for the first time in (Petroni et al., 2016). These previous studies have shown that the three strongest correlated components are usually enough to explain most of the correlation. In the technical validation section below, we have thus limited the sum to the first three components.

#### Paradigm #4 (Active): Contrast Change Detection

##### Task Overview

Our contrast change detection task is based on a recently presented EEG paradigm innovation that enables the isolation and simultaneous tracing of neural dynamics at the three major processing stages underlying simple sensorimotor decisions: sensory evidence encoding, evidence accumulation over time, and motor preparation (O’Connell et al., 2012). Here we employed a modified version of that task in order to probe fluctuations in attentional engagement in addition to these three sensory-motor processing levels.

Simple sensory-motor decision making – i.e., choosing a course of action based on a sensory judgment – can be regarded as a core component of a large portion of human behavior, and of almost any neuropsychological test administered in clinical settings. Such decisions require the momentary encoding of sensory information necessary for the decision (evidence), the sequential integration of that evidence into a “decision variable,” and the concomitant preparation of an appropriate action. Whereas typical EEG tasks involve sudden-onset, discrete stimuli that evoke a complex set of overlapping components on the scalp, only a small proportion of which relate to the relevant computations underlying task performance, our contrast change detection paradigm uses gradual-change targets, thereby eliminating transient, task-irrelevant sensory-evoked signals and thus fully unmasks the neural processes of decision formation. By asking subjects to indicate detection of a change in contrast of a continuously presented, flickering visual stimulus, an independent and continuous neurophysiological measure of the momentary sensory input to the decision process can also be extracted. In tandem, motor preparatory activity such as contralateral pre-motor movement-selective beta-band (16–30 Hz) activity can be traced (Pfurtscheller et al., 1999; Donner et al., 2009). Thus, discrete, freely evolving neural signatures of sensory evidence encoding, decision formation and motor preparation, can be isolated using this paradigm.

In the present task battery, we employ a two-alternative version of the contrast change detection paradigm, whereby, instead of detecting a change to a single stimulus component with a single response, subjects must monitor the relative contrast of two simultaneous stimuli for gradual changes and select one of two responses to indicate the direction of the change. The reasoning behind this is that fluctuations in the sensory evidence (the difference in response to the two stimuli to be compared) can be dissociated to some degree from fluctuations in general arousal or levels of sustained attention (non-selective changes common to both responses). Such fluctuations are of considerable interest in their own right, both in clinical and basic neuroscience (Dockree et al., 2005; Dockree et al., 2007; Smallwood et al., 2008; Smallwood et al., 2009), and are an inherent aspect of the change detection task which is performed continuously in long, uninterrupted blocks with infrequent and unpredictable target onsets.

##### Stimuli & Experimental Design

The contrast change detection paradigm is designed to enable isolation of the neural signatures of sensory evidence encoding, accumulation, and motor preparation without the need for complex signal processing beyond elementary epoch averaging and spectral estimation (O’Connell et al., 2012). In the present task, subjects continuously viewed an annular pattern (inner radius: 1°; outer radius 6°) composed of two overlaid gratings tilted 45° to the left and 45° to the right of vertical, which continuously phase-reversed at distinct rates of 20 Hz and 25 Hz, respectively. At baseline (in between targets), both gratings had an equal contrast of 50%. Participants were asked to maintain fixation on a point in the center of this stimulus, and to detect contrast-change targets, where one grating gradually increased to 100% and the other simultaneously decreased to 0%. They were asked to make a left-hand button click for targets in which the left-tilted grating increased in contrast, and to make a right-hand click for right-tilted increases. Twelve of each of these two target types were presented in each 3.1-minute block, in random order. The changes in contrast from 50 to 100% occurred linearly over 1600 ms, with an immediate 800 ms linear return to 50%. Beginning immediately at the end of each target, the 50% contrast baseline stimulus was presented for an inter-target interval of 2.8, 4.4 or 6 sec. Also, immediately following target end, feedback was presented in the form of a smiley (correct click) or sad face (incorrect click or no click) for the first 400 ms of the inter-target interval. If a subject missed three consecutive targets, a short voice recording was played, which saying, “You just missed three targets in a row. Please focus again.” In the current dataset, each subject completed 3 blocks of this task.

##### Participant Instructions

“Fixate on the central dot. Press the LEFT button with LEFT hand when the LEFT-tilted pattern gets stronger. Press the RIGHT button with RIGHT hand when the RIGHT-tilted pattern gets stronger. Work as quickly as you can without making mistakes. Press the mouse button to begin.”

##### Output Measures

By design, the principal components of activity on this task are the SSVEP over occipital scalp sites, the event-related potential over centro-parietal scalp sites, and decreases in Mu (8–13 Hz) and Beta (16–30 Hz) spectral amplitude over left/right motor cortical areas (C3/C4), which reflect sensory evidence encoding, evidence accumulation and motor preparation, respectively (O’Connell et al., 2012). Each of these signals has been shown to bear a systematic relationship with the timing and accuracy of the participant’s detection responses. Since this task version involves two-alternative decisions mapped to the left and right hands, the relative preparation for the two alternative actions can also be tracked via the lateralized readiness potential derived by subtracting ERP traces from motor cortical sites of the two hemispheres (Gratton et al., 1988; O’Connell et al., 2012). In addition to these measures, posterior parietal alpha-band activity can be analyzed to provide measures of vigilant attentional state. In principle, because the monitoring task is performed continuously and stimulation is continuous, neural activity measures are potentially informative on cognitive/perceptual states and processes at any point during the block of task performance.

#### Paradigm #5 (Active): Sequence Learning

##### Task Overview

In order to evaluate the neural correlates of declarative learning, we included an explicit visual sequence learning paradigm, in which subjects repeatedly view a fixed sequence of flashed visual locations and attempt to memorize it in order to make regular intermediate recall reports. This task was originally developed by Moisello, Ghilardi and colleagues as a control condition for the examination of spectral EEG signatures of visuo-motor learning (Moisello et al., 2013), and was recently shown to be highly informative in its own right, in providing reliable indices of memory formation and surprise-modulated stimulus processing that related systematically to the ongoing progress of learning (Steinemann et al., 2016). An important aspect of the paradigm is that the information to be remembered (flashed location) is of the most elementary kind and computed very rapidly in the brain, so that perceptual decisions regarding the immediately presented item are completed quickly, allowing the longer-lasting neural signatures of memory formation to be reliably distinguished from the short-lived processes of immediate stimulus identification.

During the task, participants were asked to observe and memorize a single sequence of elements over repeated observations. This enables the possibility to track the progress of gradual memory formation through regular behavioral recall, as an individual element goes from being completely unknown to fully committed to memory. Rather than making comparisons among different complex items as is commonly done in the field (Schacter and Wagner, 1999; Wagner et al., 1999), which may differ in sensory characteristics and/or semantic content, this paradigm enables comparisons across successive learning states for each of a set of uniform, highly reduced, and semantically unloaded stimuli. This neural and behavioral tracking of the gradual learning progress in a way that cannot be done using typically employed paradigms with dichotomous subsequent recall outcomes (remembered vs. forgotten) (Karis et al., 1984; Paller et al., 1987; Neville et al., 1996).

##### Stimuli & Experimental Design

In the current task battery we employed an adapted version of the task of Steinemann et al. (2016). Participants were asked to view a sequence of 10 flashed-circle stimuli, which appeared among 8 possible, marked locations on the screen. The same sequence was presented a total of 5 times; after viewing each presentation, the participant attempted to reproduce the sequence to the best of their ability by sequentially clicking the different locations using a computer mouse. In pilot testing, we observed a floor effect on this 10-item sequence version in children younger than 9 years old; therefore, in the present study, participants 8 years and below were shown a shorter sequence of 8 items displayed among 6 possible locations. There was no restriction on the time provided to report the recalled sequence, and no feedback was provided throughout the task. Visual stimuli consisted of filled white circles with a diameter of 1 cm presented at eight different equidistant spatial locations on a radius of 5 cm eccentricity, and were continuously marked by static circular outlines (see Fig. 1 in (Steinemann et al., 2016). Stimuli were presented (and gradually faded out) for 200 ms, with an inter-stimulus interval of 1300 ms. Throughout the task, subjects were asked to hold eye fixation on a central fixation point (yellow dot). Before the main task recording, a training block was administered, consisting of 5 stimuli on the same 8 locations, in order to familiarize the subjects with the tasks and to confirm their comprehension of them. Feedback was provided for the training task only. The duration of this paradigm varied between 8–15 min, depending on the speed of recall reports.

##### Participant Instructions

“Fixate on the yellow dot. Try to remember the sequence of the flashing dots. The SAME sequence will be repeated 5 times. After each round you have to give a response. If you do not know all the locations guess the others. Press the mouse button to begin.”

##### Output Measures

In the approach of Steinemann et al. (2016), trials were categorized as “still-unknown”, “newly-learned” or “known” based on the participants’ recall reports, and the average ERPs for these learning states were directly compared to examine processes of immediate stimulus identification and their modulation by “surprise,” which reduced over the course of learning, and processes of memory formation which were especially strong at the point where a given item was newly learned. For the purposes of the current paper, we analyzed behavioral recall performance as well as these neural correlates over the successive blocks of sequence observation, which provides a simpler, but related, view on the progress of learning over the task. The process of immediate stimulus identification is reflected in a “P300” component measured over centro-parietal sites. The P300 is a centro-parietal positivity occurring roughly 300 ms or later after stimulus onset, which famously indexes the level of “surprise”, i.e., the degree to which a stimulus was unexpected (Donchin, 1981; Mars et al., 2008). Recently it has been established that the P300 corresponds to the centro-parietal positivity (CPP), which reflects the accumulation of evidence for a decision, and it has been suggested that its sensitivity to surprise may arise from the setting of higher accumulation thresholds for unexpected stimuli (O’Connell et al., 2012; Twomey et al., 2015). In the sequence learning paradigm, as learning progresses, the location of the stimuli becomes increasingly less surprising, and therefore P300 amplitude decreases systematically. In fact, the degree of P300 reduction from the first to second block of sequence observation was found to correlate significantly with behavioral measures of the speed of learning (Steinemann et al., 2016), highlighting the potential value of such measures.

#### Paradigm #6 (Active): Symbol Search

##### Task Overview

As our final, “active and complex” paradigm, we chose to emulate a standard neuropsychological test in widespread, routine clinical use for assessing “processing speed” in children. We chose the particular construct of processing speed because it is a good example among a wide range of clinical metrics that are almost universally employed yet imprecisely defined, with many conceivable computational explanations that can account for variation in the lumped, unitary score that is ultimately recorded on completion of the test. The “processing speed” construct has been defined as the ability to focus attention, quickly scan, and discriminate between (visual) information, and is known to be sensitive to factors such as motivation, difficulty working under time pressure, and motor coordination (Wechsler, 2004). Previous studies have associated processing speed with age, reading performance, and psychiatric and neurological disorders (Donders et al., 2001; Salthouse and Ferrer-Caja, 2003; Eckert, 2011; Duering et al., 2014). We selected a test of processing speed in the current dataset due to the obvious scope for using neurophysiological and eye tracking measures to deconstruct performance into a richer set of computationally tractable component processes.

The specific paradigm used here was a computerized version of the Symbol Search subtest of the Wechsler Intelligence Scale for Children IV (WISC-IV), which together with the subtests Coding and Cancellation makes up the Process Speed Index (PSI) (Burgess et al., 1992; Lezak, 1995; Wechsler, 2004). The Symbol Search subtest is designed to assess the speed and accuracy with which a child can process nonverbal information. High scores require rapid and accurate processing of visual symbols that have no *a priori* meaning, which hinges on processing efficiency at several levels including motor, cognitive, and decisional and memory processes (e.g. Royer et al., 1981; Joy et al., 2004). For example, a participant needs to (a) detect and encode the target symbols; (b) hold this information in short-term and/or working memory; (c) process each of the symbols in the search set, whether in turn or in parallel to some degree; (d) identify the symbol among the search set that matches one of the target symbols, or conclude that there is no match; (e) select and initiate the appropriate response. This paradigm further enables the study of different strategies or performance styles that might cause a decreased performance, such as excessive carefulness (i.e., double-checking, or ‘making sure’).

It is not entirely clear which components of symbol search task performance are affected by decreases in processing speed, as the standard application of the task provides only one overall behavioral score (number relatively correct); little or no information on the underlying etiology of low performance is offered. Our on-line simultaneous acquisition of eye tracking and EEG data during this test thus stands to provide substantial further insights. We believe this integrated EEG/eye tracking approach will allow us to decompose the processing speed task into interpretable components of cognitive and perceptual processing, such as working memory, distractibility, uncertainty, and sustained attention.

##### Stimuli & Experimental Design

The visual geometric stimuli consisted of black symbols with a size of 1cm width and 1cm height (Figure 7A). As on each page of the paper version, 15 trials were presented at a time on the screen. Each row contained two target symbols and five search symbols, arranged horizontally across the row. Participants were instructed to indicate for each row, by mouse-click (mark either the yes or no checkbox), whether either of the target symbols matched with any of the five search symbols. The participants had the option to correct their initial responses if they desired. Participants were instructed to solve as many rows, or trials, as possible within two minutes. Before beginning the actual paradigm, participants performed a training block with 4 trials, for which they received feedback, to ensure their comprehension of the task. No feedback was provided throughout the actual task.

Once a participant finished all 15 trials, they pressed the “next page” button to advance onward. There were 4 pages (a maximum of 60 trials) in total. No participant ever reached the end of the 60 trials.

##### Participant Instructions

“The task is to figure out if either one of the two first symbols are presented again in the same line. Press with the left mouse button YES and NO boxes to select your answer. If you accidently press the wrong button you can make a correction by simply clicking on the other response. You have 2 minutes to solve as many trials as possible.”

##### Output Measures

In contrast to the traditional pen and paper administration of the symbol search task, our computerized, multimodal approach allows for the generation of a range of measures rather than a single summary score. These included, but were not limited to: time spent looking at each symbol, the number of saccade steps, number of repetitions, pupil size, and the protracted gaze dwell times for each sub-region of the screen. These measures supply additional information on participants’ strategies for completing the task, and on why they might do well or poorly. This eye tracking data can further be complemented with topographic spatial and power analyses of the concurrently acquired EEG data.

### EEG and Eye Tracking Preprocessing Steps

#### EEG Data Extraction

The data shared in this project are available as raw data, but also preprocessed. The MATLAB code for the preprocessing can be found at https://github.com/amirrezaw/automagic. The data from each paradigm is saved as a separate file. In the first step of preprocessing, EEG data were imported in MATLAB *(pop_readegi.m)* and the triggers and latencies for each paradigm were extracted. Based on this information, the specific order and availability/existence were determined for each paradigm and participant. The electrodes in the outermost circumferences (chin and neck) were excluded to a standard 111-channel electrode array (Perrin et al., 1987).

#### Electrode Quality Check

Bad electrodes were identified and replaced. Identification of bad electrodes was based on probability, kurtosis, and frequency spectrum distribution of all electrodes. A channel was defined as a bad electrode when recorded data from that electrode had a variance more than 3 standard deviations away from the mean across all other electrodes. This was realized with the eeglab MATLAB function: “pop_rejchan.m”. Subsequently bad electrodes were interpolated by using a using spherical spline interpolation (Perrin et al., 1989, 1990) ‘eeg_interp.m’. Moreover, after automatic scanning, noisy channels were selected by visual inspection and interpolated or replaced entirely by zeros (for the calculation of the ISC measures to eliminate the channel’s contribution in subsequent calculation of covariance matrices).

#### Artifact Signal Correction

One hundred and nine EEG channels were used for scalp recordings, while 6 EOG channels were used for artifact removal. The rest of the channels lying mainly on the neck and face were discarded before data analysis. Data were then high-pass filtered at 0.1 Hz and notch filtered at 59–61 Hz. Eye artifacts were removed by linearly regressing the EOG channels from the scalp EEG channels. Next, a robust Principal Components Analysis (PCA) algorithm, the inexact Augmented Lagrange Multipliers Method (ALM, (Lin et al., 2010) removed sparse noise from the data. Briefly, the ALM recovers a low-rank matrix, A, efficiently and accurately from a corrupted data matrix D = A + E, where some entries of the additive errors E may be arbitrarily large. Finally, the entire dataset for each subject was visually inspected in order to discard whole block and/or paradigm recordings that remained noisy after the automatic and manual noise removal methods. All signal processing was performed offline using MATLAB software (MathWorks, Natick, MA, USA).

#### Eye Tracking Data Extraction

Saccades and fixations were detected with an adaptive velocity-based algorithm. Briefly, a blink can be regarded as a special case of a fixation, where the pupil diameter is either zero or outside a dynamically computed valid pupil, or the horizontal and vertical gaze positions are zero. The algorithm identifies fixations as groups of consecutive points within a particular dispersion. It uses a moving window that spans consecutive data points checking for potential fixations. The moving window begins at the start of the protocol and initially spans a minimum number of points, determined by the given Minimum Fixation Duration (here: 50 ms) and sampling frequency. The algorithm then checks the dispersion of the points in the window by summing the differences between the points’ maximum and minimum x and y values and comparing that to the Maximum Dispersion Value; so if [max(x) – min(x)] + [max(y) -min(y)] > Maximum Dispersion Value, the window does not represent a fixation, and the window moves one point to the right. If the dispersion is below the Maximum Dispersion Value (here: 50 pixels), the window represents a fixation. In this case, the window is expanded to the right until the window’s dispersion is above threshold. The final window is registered as a fixation at the centroid of the window points with the given onset time and duration. Following this process, a saccade event is created between the newly and the previously created blink or fixation.

### Code availability

The codes for the EEG preprocessing can be found here: https://github.com/amirrezaw/automagic. Code for the ISC analysis is available here: http://parralab.org/isc. All the analyses were performed with MATLAB 2014a and EEGlab 13.3.2b.

### Data Records

#### Data privacy

All data are de-identified and participants gave permission for their data to be openly shared as part of the informed consent process.

#### Distribution for use

Raw and preprocessed EEG data, as well as eye tracking data can be accessed through the 1000 Functional Connectomes Project and its International Neuroimaging Data-sharing Initiative (FCP/INDI) based at www.nitrc.org (http://fcon_1000.projects.nitrc.org/indi/cmi_eeg/). EEG data are available openly, along with basic phenotypic data (age, sex, handedness, completion status of EEG paradigms, and known diagnosis status) and performance measures for the EEG paradigms. Public data are distributed under the Creative Commons, Attribution NonCommercial Share Alike License (https://creativecommons.org/licenses/by-nc-sa/3.0/).

The more extensive phenotypic data (e.g., behavioral questionnaires, abbreviated intelligence and achievement testing; see Table 3) may be accessed through the Collaborative Informatics and Neuroimaging Suite (COINS) Data Exchange (http://coins.mrn.org/dx). These data are protected by a Data Usage Agreement (DUA), which investigators must complete and have signed by an authorized institutional official before receiving access (the DUA can be found at: http://fcon_1000.projects.nitrc.org/indi/cmi_eeg/phenotypic.html). The DUA is based upon that of the NKI-Rockland Sample, which does not attempt to restrict or curate the focus of analyses, but does require users to agree not to attempt re-identification of participants under any circumstances.

**Table 3.**
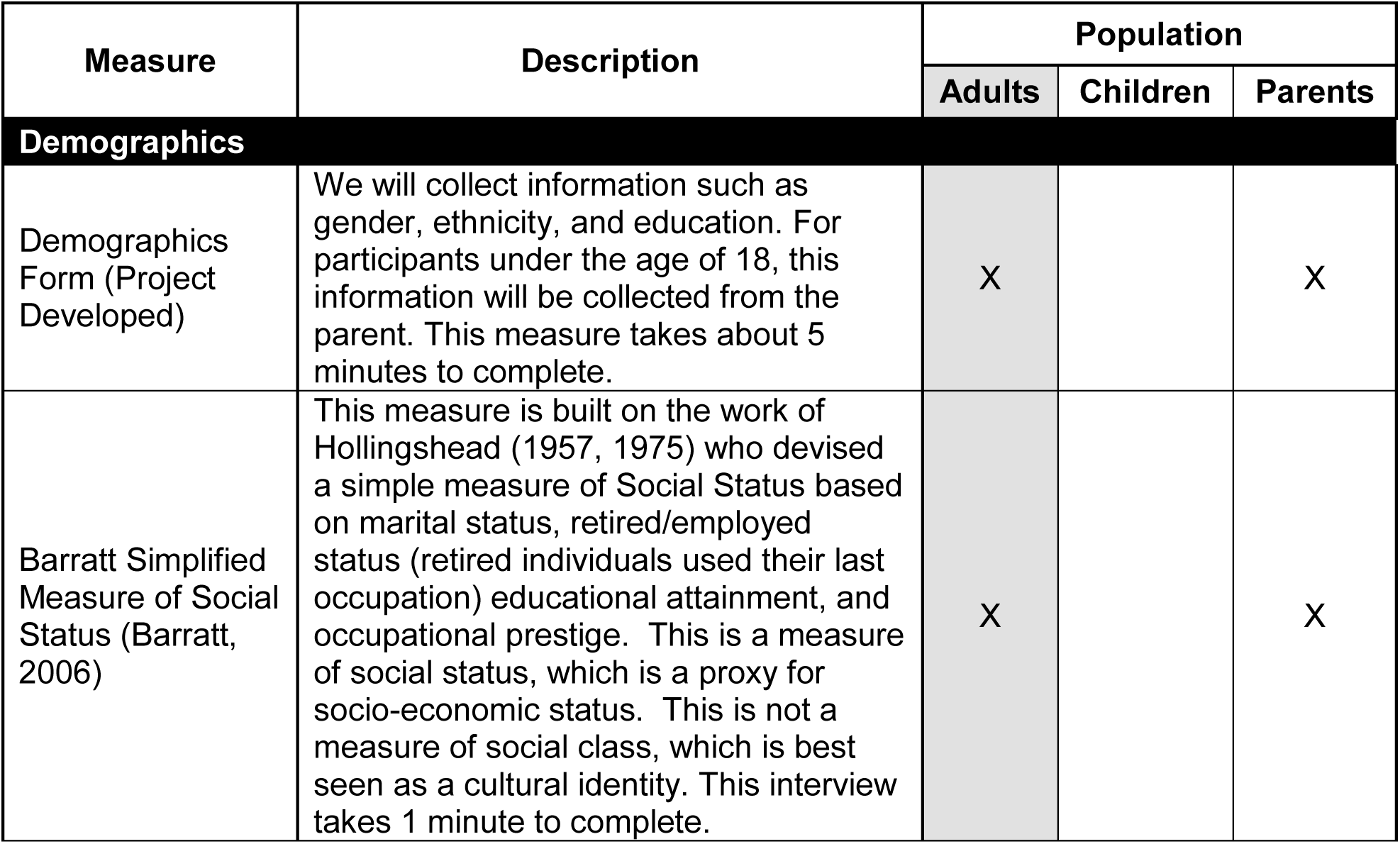
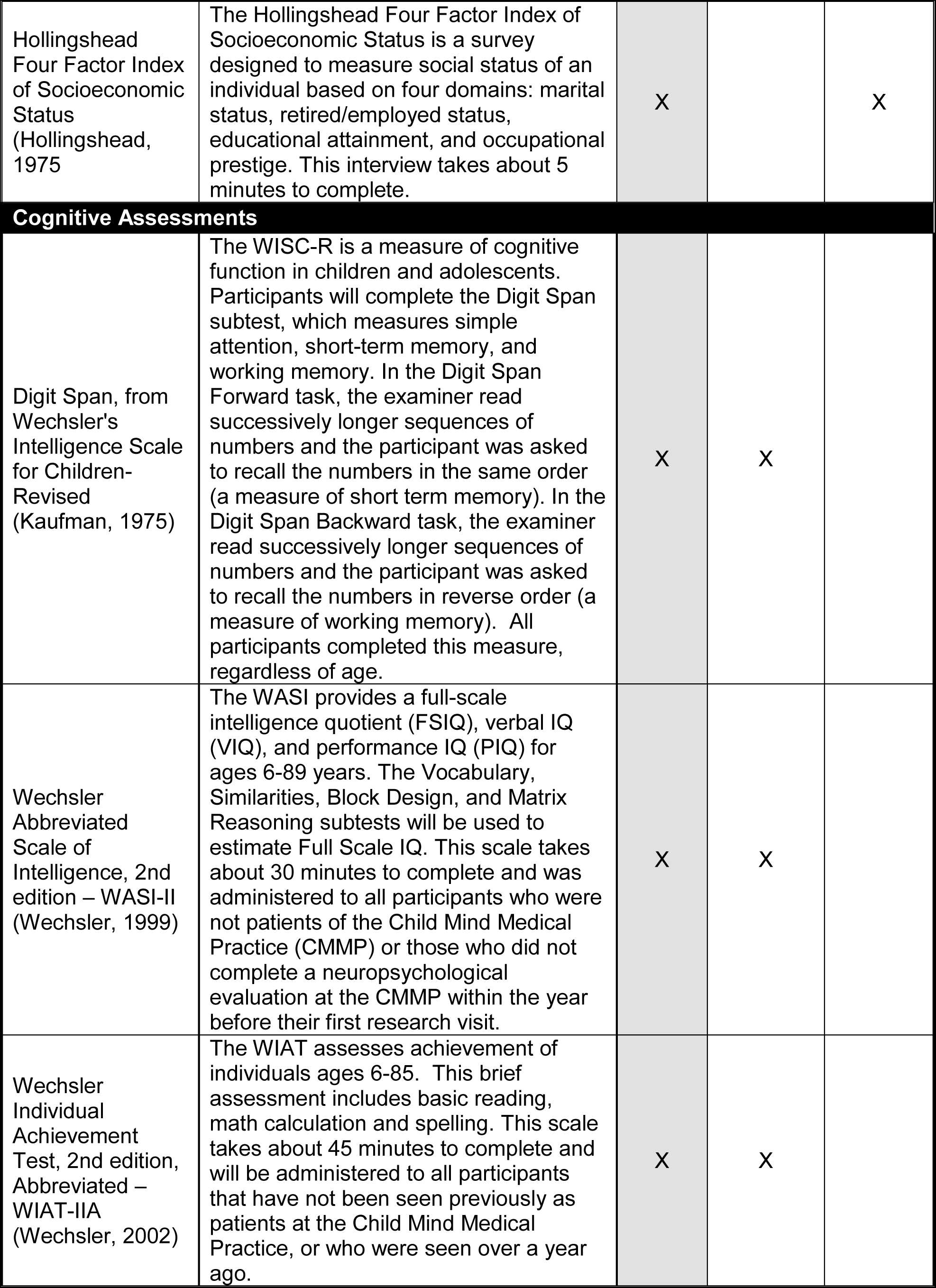
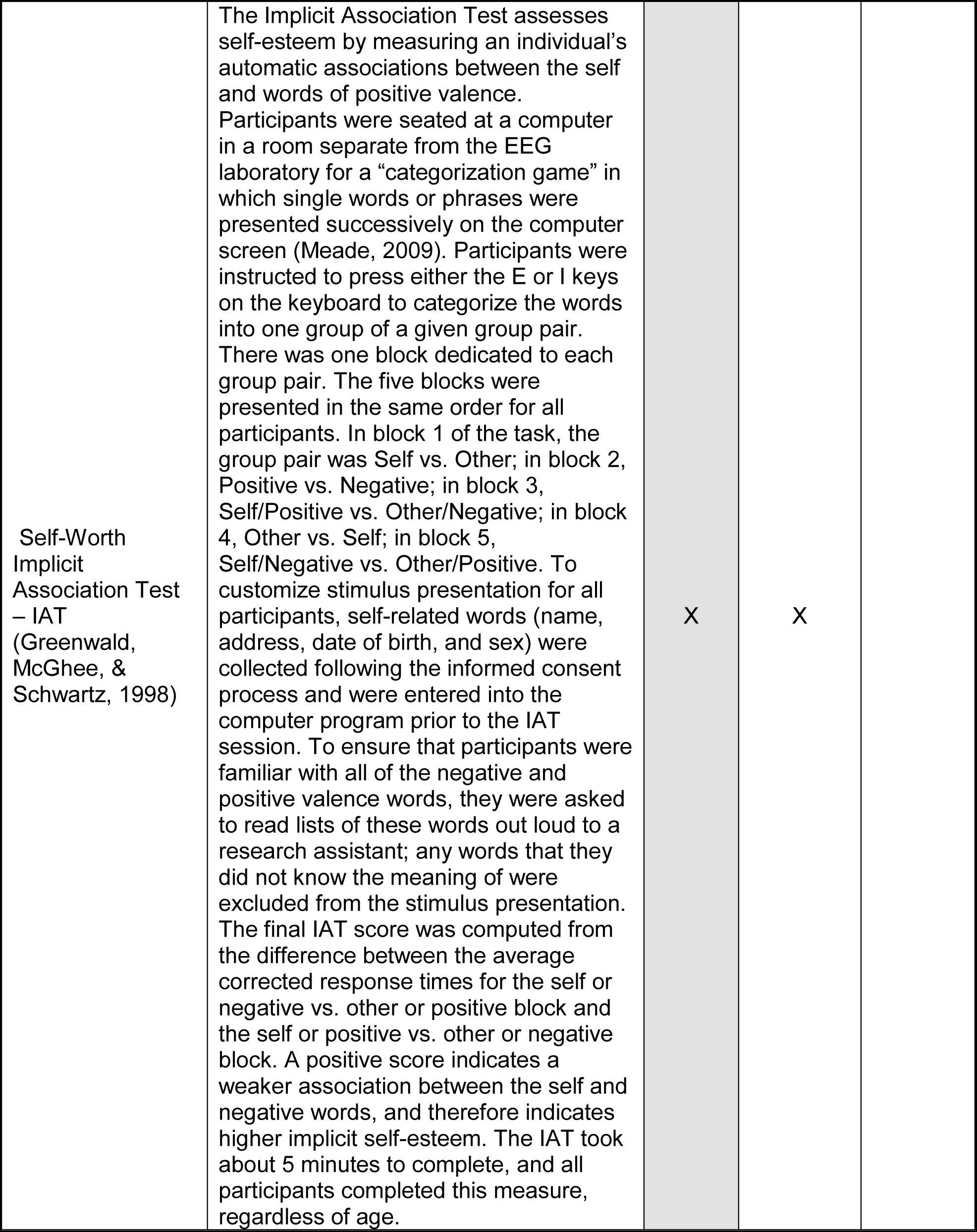
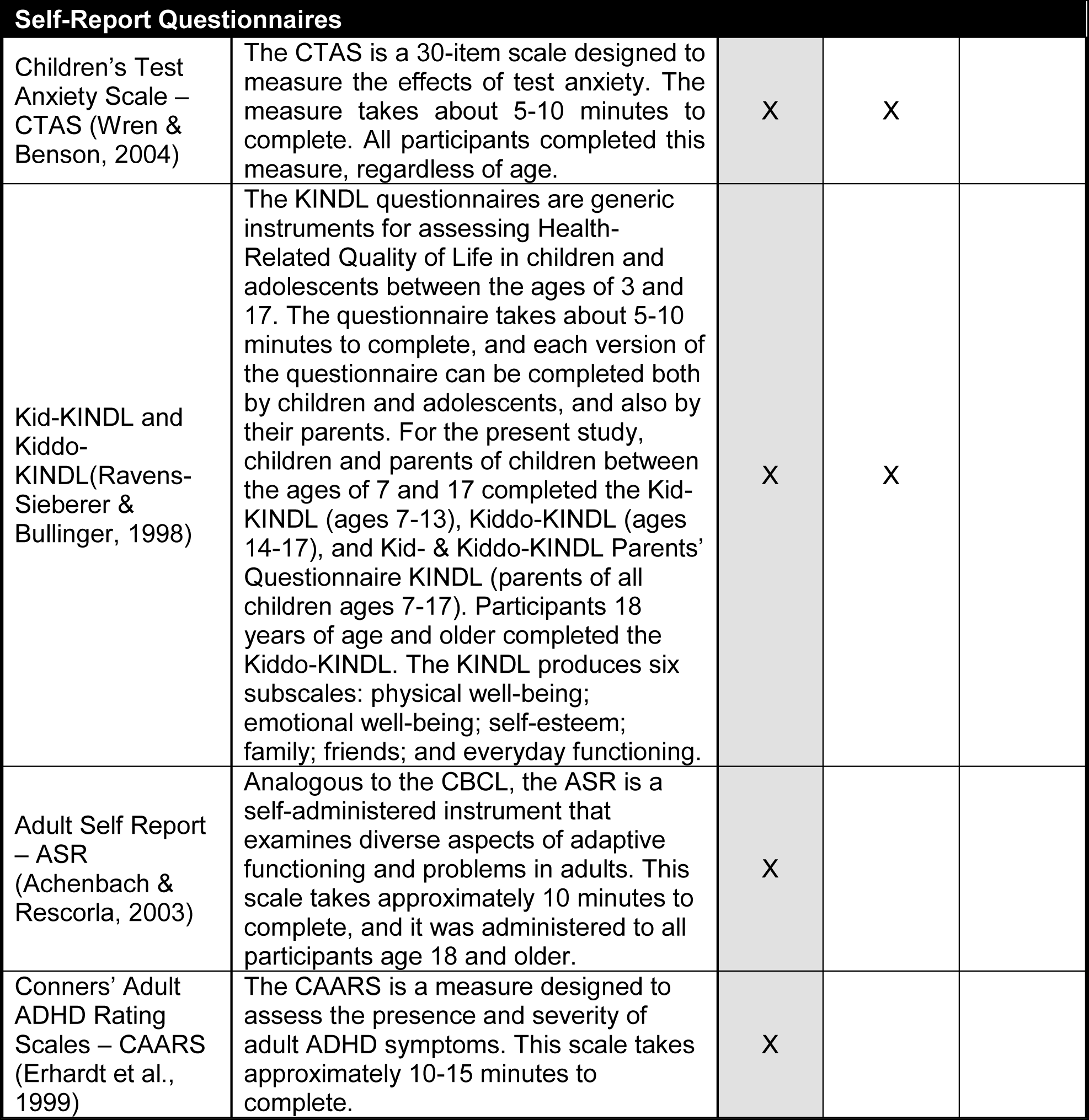
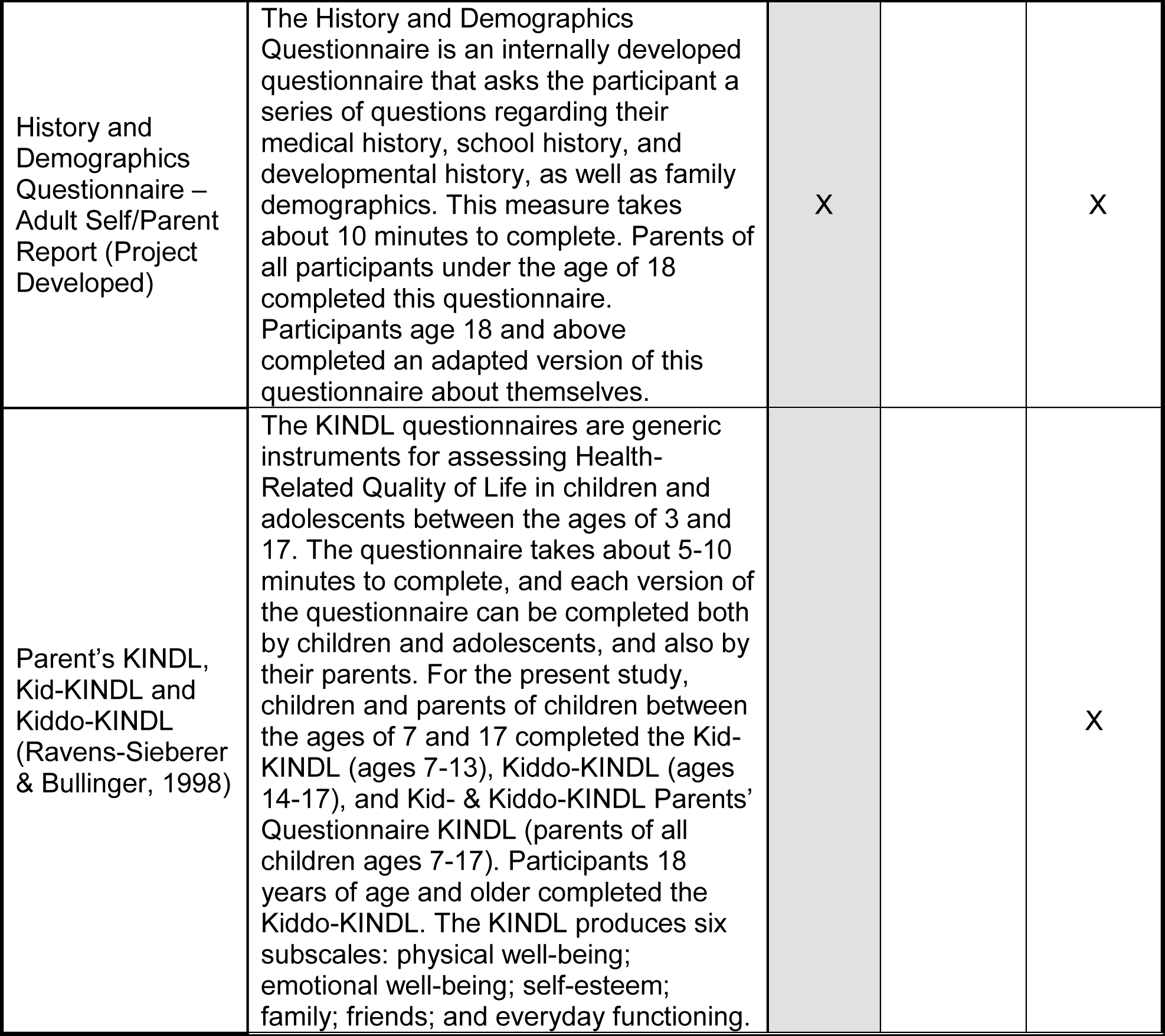
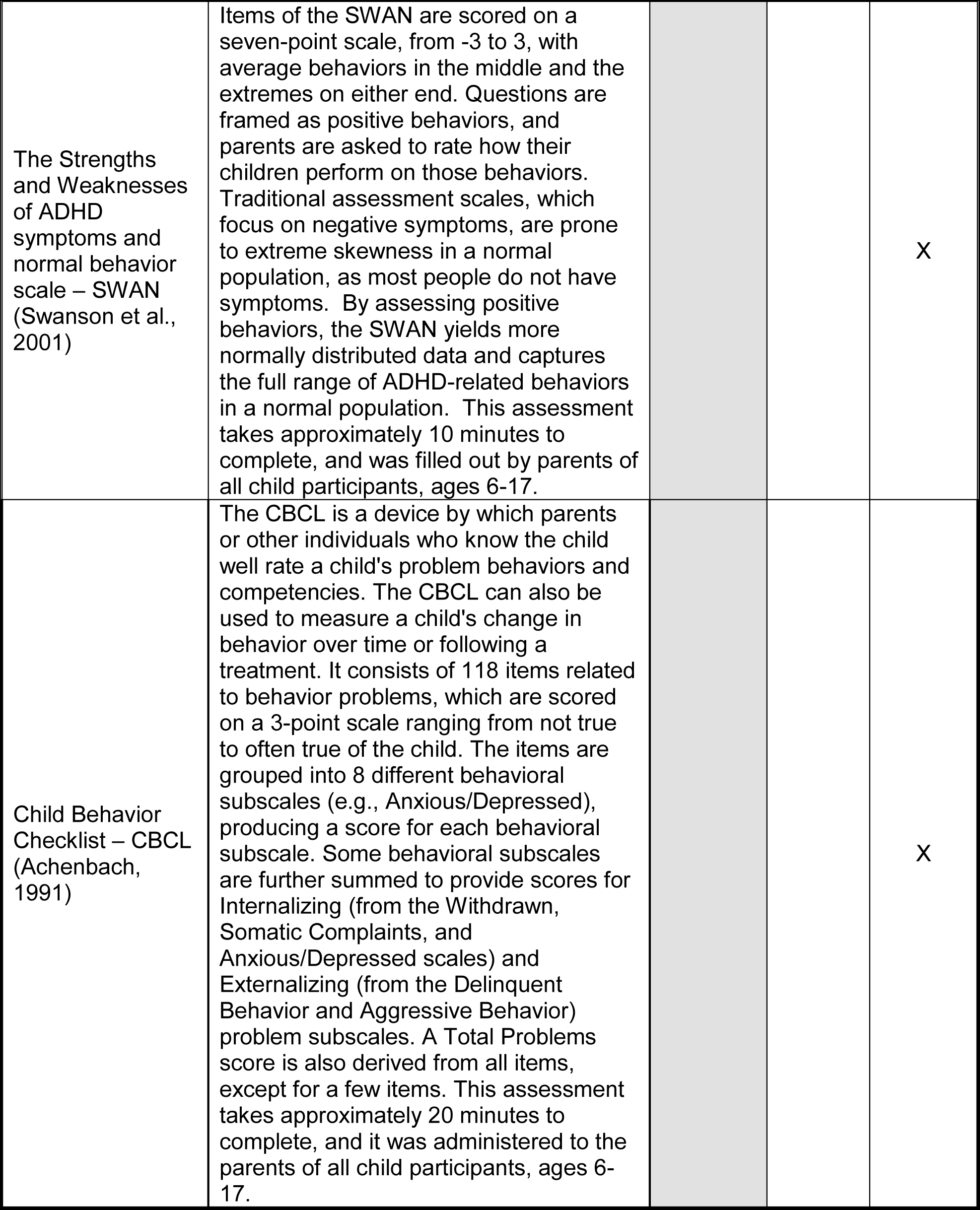
Phenotypic Data Available. The complete list of the phenotypic data for each subjects is listed. The name of the cognitive test or questionnaire, a description of the measure, and target subjects are described.

| Measure | Description | Population |  |  |
| --- | --- | --- | --- | --- |
|  |  | Adults | Children | Parents |
| Demographics |  |  |  |  |
| Demographics Form (Project Developed) | We will collect information such as gender, ethnicity, and education. For participants under the age of 18, this information will be collected from the parent. This measure takes about 5 minutes to complete. | X |  | X |
| Barratt Simplified Measure of Social Status (Barratt, 2006) | This measure is built on the work of Hollingshead (1957, 1975) who devised a simple measure of Social Status based on marital status, retired/employed status (retired individuals used their last occupation) educational attainment, and occupational prestige. This is a measure of social status, which is a proxy for socio-economic status. This is not a measure of social class, which is best seen as a cultural identity. This interview takes 1 minute to complete. | X |  | X |
| Hollingshead Four Factor Index of Socioeconomic Status (Hollingshead, 1975) | The Hollingshead Four Factor Index of Socioeconomic Status is a survey designed to measure social status of an individual based on four domains: marital status, retired/employed status, educational attainment, and occupational prestige. This interview takes about 5 minutes to complete. | X |  | X |
| **Cognitive Assessments** |  |  |  |  |
| Digit Span, from Wechsler's Intelligence Scale for Children-Revised (Kaufman, 1975) | The WISC-R is a measure of cognitive function in children and adolescents. Participants will complete the Digit Span subtest, which measures simple attention, short-term memory, and working memory. In the Digit Span Forward task, the examiner read successively longer sequences of numbers and the participant was asked to recall the numbers in the same order (a measure of short term memory). In the Digit Span Backward task, the examiner read successively longer sequences of numbers and the participant was asked to recall the numbers in reverse order (a measure of working memory). All participants completed this measure, regardless of age. | X | X |  |
| Wechsler Abbreviated Scale of Intelligence, 2nd edition – WASI-II (Wechsler, 1999) | The WASI provides a full-scale intelligence quotient (FSIQ), verbal IQ (VIQ), and performance IQ (PIQ) for ages 6-89 years. The Vocabulary, Similarities, Block Design, and Matrix Reasoning subtests will be used to estimate Full Scale IQ. This scale takes about 30 minutes to complete and was administered to all participants who were not patients of the Child Mind Medical Practice (CMMP) or those who did not complete a neuropsychological evaluation at the CMMP within the year before their first research visit. | X | X |  |
| Wechsler Individual Achievement Test, 2nd edition, Abbreviated – WIAT-IIA (Wechsler, 2002) | The WIAT assesses achievement of individuals ages 6-85. This brief assessment includes basic reading, math calculation and spelling. This scale takes about 45 minutes to complete and will be administered to all participants that have not been seen previously as patients at the Child Mind Medical Practice, or who were seen over a year ago. | X | X |  |
| Self-Worth<br>Implicit<br>Association Test<br>– IAT<br>(Greenwald,<br>McGhee, &<br>Schwartz, 1998) | The Implicit Association Test assesses self-esteem by measuring an individual's automatic associations between the self and words of positive valence. Participants were seated at a computer in a room separate from the EEG laboratory for a “categorization game” in which single words or phrases were presented successively on the computer screen (Meade, 2009). Participants were instructed to press either the E or I keys on the keyboard to categorize the words into one group of a given group pair. There was one block dedicated to each group pair. The five blocks were presented in the same order for all participants. In block 1 of the task, the group pair was Self vs. Other; in block 2, Positive vs. Negative; in block 3, Self/Positive vs. Other/Negative; in block 4, Other vs. Self; in block 5, Self/Negative vs. Other/Positive. To customize stimulus presentation for all participants, self-related words (name, address, date of birth, and sex) were collected following the informed consent process and were entered into the computer program prior to the IAT session. To ensure that participants were familiar with all of the negative and positive valence words, they were asked to read lists of these words out loud to a research assistant; any words that they did not know the meaning of were excluded from the stimulus presentation. The final IAT score was computed from the difference between the average corrected response times for the self or negative vs. other or positive block and the self or positive vs. other or negative block. A positive score indicates a weaker association between the self and negative words, and therefore indicates higher implicit self-esteem. The IAT took about 5 minutes to complete, and all participants completed this measure, regardless of age. | X | X |  |

| Self-Report Questionnaires |  |  |  |
| --- | --- | --- | --- |
| Children's Test Anxiety Scale – CTAS (Wren & Benson, 2004) | The CTAS is a 30-item scale designed to measure the effects of test anxiety. The measure takes about 5-10 minutes to complete. All participants completed this measure, regardless of age. | X | X |
| Kid-KINDL and Kiddo-KINDL (Ravens-Sieberer & Bullinger, 1998) | The KINDL questionnaires are generic instruments for assessing Health-Related Quality of Life in children and adolescents between the ages of 3 and 17. The questionnaire takes about 5-10 minutes to complete, and each version of the questionnaire can be completed both by children and adolescents, and also by their parents. For the present study, children and parents of children between the ages of 7 and 17 completed the Kid-KINDL (ages 7-13), Kiddo-KINDL (ages 14-17), and Kid- & Kiddo-KINDL Parents' Questionnaire KINDL (parents of all children ages 7-17). Participants 18 years of age and older completed the Kiddo-KINDL. The KINDL produces six subscales: physical well-being; emotional well-being; self-esteem; family; friends; and everyday functioning. | X | X |
| Adult Self Report – ASR (Achenbach & Rescorla, 2003) | Analogous to the CBCL, the ASR is a self-administered instrument that examines diverse aspects of adaptive functioning and problems in adults. This scale takes approximately 10 minutes to complete, and it was administered to all participants age 18 and older. | X |  |
| Conners' Adult ADHD Rating Scales – CAARS (Erhardt et al., 1999) | The CAARS is a measure designed to assess the presence and severity of adult ADHD symptoms. This scale takes approximately 10-15 minutes to complete. | X |  |
| History and Demographics Questionnaire – Adult Self/Parent Report (Project Developed) | The History and Demographics Questionnaire is an internally developed questionnaire that asks the participant a series of questions regarding their medical history, school history, and developmental history, as well as family demographics. This measure takes about 10 minutes to complete. Parents of all participants under the age of 18 completed this questionnaire. Participants age 18 and above completed an adapted version of this questionnaire about themselves. | X |  | X |
| Parent's KINDL, Kid-KINDL and Kiddo-KINDL (Ravens-Sieberger & Bullinger, 1998) | The KINDL questionnaires are generic instruments for assessing Health-Related Quality of Life in children and adolescents between the ages of 3 and 17. The questionnaire takes about 5-10 minutes to complete, and each version of the questionnaire can be completed both by children and adolescents, and also by their parents. For the present study, children and parents of children between the ages of 7 and 17 completed the Kid-KINDL (ages 7-13), Kiddo-KINDL (ages 14-17), and Kid- & Kiddo-KINDL Parents' Questionnaire KINDL (parents of all children ages 7-17). Participants 18 years of age and older completed the Kiddo-KINDL. The KINDL produces six subscales: physical well-being; emotional well-being; self-esteem; family; friends; and everyday functioning. |  |  | X |
| The Strengths and Weaknesses of ADHD symptoms and normal behavior scale – SWAN (Swanson et al., 2001) | Items of the SWAN are scored on a seven-point scale, from -3 to 3, with average behaviors in the middle and the extremes on either end. Questions are framed as positive behaviors, and parents are asked to rate how their children perform on those behaviors. Traditional assessment scales, which focus on negative symptoms, are prone to extreme skewness in a normal population, as most people do not have symptoms. By assessing positive behaviors, the SWAN yields more normally distributed data and captures the full range of ADHD-related behaviors in a normal population. This assessment takes approximately 10 minutes to complete, and was filled out by parents of all child participants, ages 6-17. |  |  | X |
| Child Behavior Checklist – CBCL (Achenbach, 1991) | The CBCL is a device by which parents or other individuals who know the child well rate a child's problem behaviors and competencies. The CBCL can also be used to measure a child's change in behavior over time or following a treatment. It consists of 118 items related to behavior problems, which are scored on a 3-point scale ranging from not true to often true of the child. The items are grouped into 8 different behavioral subscales (e.g., Anxious/Depressed), producing a score for each behavioral subscale. Some behavioral subscales are further summed to provide scores for Internalizing (from the Withdrawn, Somatic Complaints, and Anxious/Depressed scales) and Externalizing (from the Delinquent Behavior and Aggressive Behavior) problem subscales. A Total Problems score is also derived from all items, except for a few items. This assessment takes approximately 20 minutes to complete, and it was administered to the parents of all child participants, ages 6-17. |  |  | X |

#### EEG data organization

The data on the AWS site are stored in folders by participant. Each participant’s folder contains one EEG folder, one eye tracking folder and one behavioral folder. In the EEG folder the EEG data are available as of the simple binary file (<ID number>.raw) and ascii file (<ID number>.csv) for each paradigm. There are also two eye tracker files for each paradigm: one file is segmented into blinks, saccades and fixations (<ID number>.txt). The other file is an unsegmented file (<ID number>.txt), with the eye tracking information for each sample. Furthermore, a behavioral folder contains a MATLAB file (<ID number>.mat) for each paradigm, which includes the information about the paradigm itself, including: inter-trial interval, triggers, number of trials, response selection and reaction time (if available). For users who intend to use the data without MATLAB, this information is also available as.csv files. Each subject’s folder requires on average 5GB storage space. A “MIPDB_EEG_Readme” folder contains “Readme” files for each paradigm about the variables and paradigm parameters.

## Technical Validation

### Resting EEG

Following standardized EEG preprocessing (described in the Methods section), the data were filtered between 1.5 and 30 Hz and segmented into eyes-closed and eyes-open segments. Only the eyes-closed segments were further analyzed for display here. The artifact-free EEG was recomputed against the average reference and segmented into 2-second epochs. In a second step, a discrete Fourier transformation algorithm was applied to the 2-second epochs. In resting-state EEG, the spectral amplitude of the signals is typically assumed to be of interest; therefore the power spectrum of 1.5–30 Hz (resolution: 0.5 Hz) was calculated. The spectra for each channel were averaged over all epochs for each subject. Next, the group mean spectral amplitude was computed and displayed as an average over all electrodes (Figure 2A) and for each electrode individually (Figure 2B). Finally, the group mean relative power spectra data were integrated for the following 7 frequency bands classification proposed by (Kubicki et al., 1979): delta (1.5–3.5 Hz); theta (4–8 Hz); alpha 1 (8.5–10 Hz); alpha 2 (10.5–12 Hz); beta 1 (12.5–18 Hz); beta 2 (18.5–21 Hz); beta 3 (21.5–30 Hz) as well as the theta / (beta1+2) ratio, which is often used in ADHD research (Arns et al., 2013) (Figure 2C). As can be seen from the figure, the expected distribution of spectral amplitudes in resting-state EEG data was obtained (Chen et al., 2008; Barry et al., 2009). These spectral measures have been sensitive and successful for describing, for instance, age-related EEG changes or various clinical conditions of developmental disorders (Michel et al., 2009).

**Figure 2.**
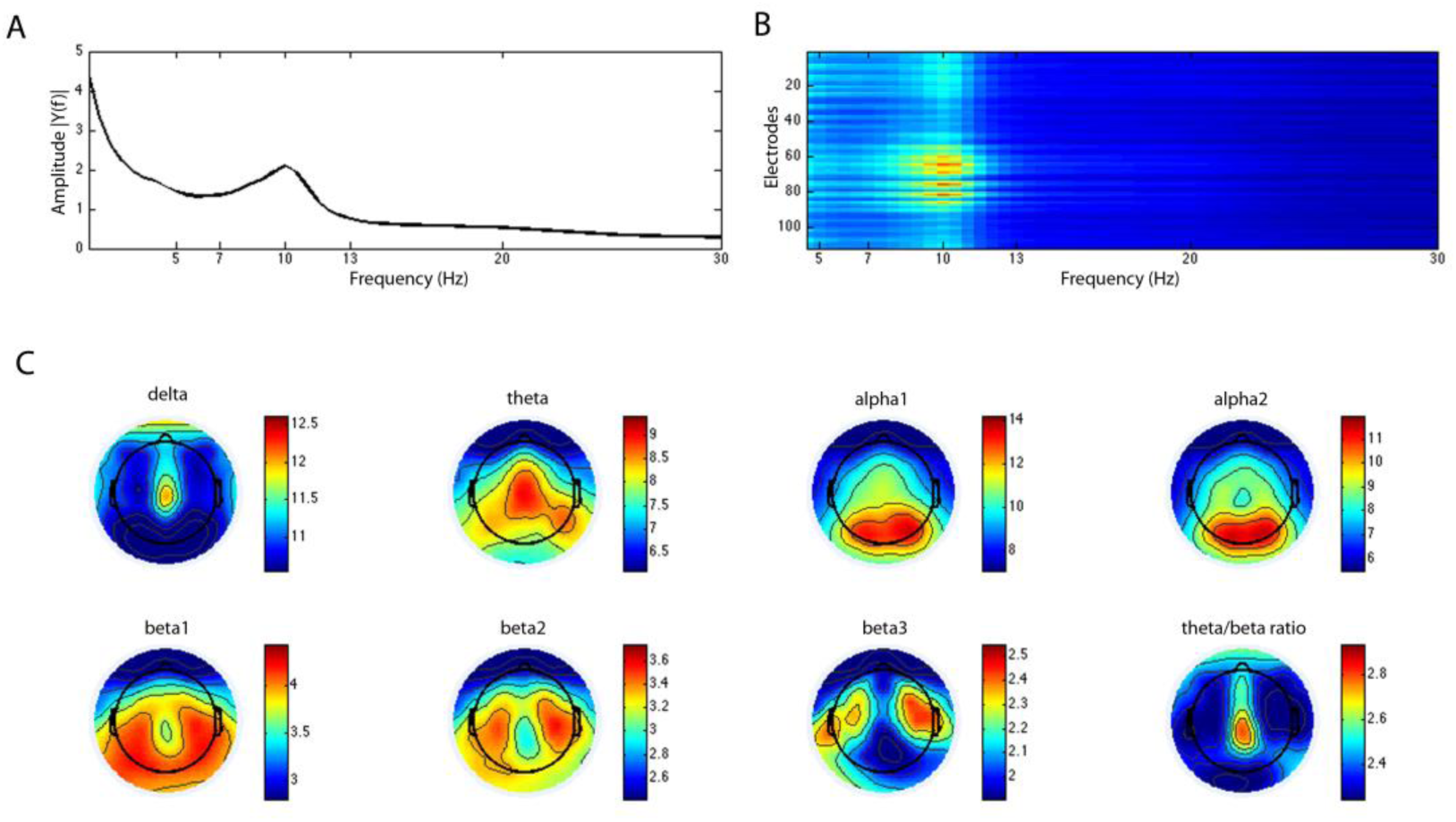
Resting EEG. The spectral amplitude was averaged over all subjects and displayed as a mean over all electrodes (A) and for each electrode individually (B). Figure C shows the topographical distribution of the group mean relative power spectra data for the different frequency bands as well as the theta / (beta1+2) ratio.

### Surround Suppression Paradigm

Following standard preprocessing, the EEG data were segmented based on the flickering stimulus onset. The segmentation was conducted for each of the 4 foreground contrasts (0% 30%, 60% and 100%) and three different background conditions (parallel, orthogonal, no background) individually. For each participant, the data were merged across the two blocks. For the purpose of technical validation, we computed 1) the SSVEP signal without background, and 2) the SSVEP across all conditions with a background. In a next step, the FFT was computed to obtain a measure of SSVEP power at 25 Hz flickering frequency. In Figure 3, we subtracted the SSVEP power for 25 Hz from its neighboring frequency bin to extract the actual evoked activity by the stimulus presentation. An in-house algorithm detected the electrode with the highest SSVEP amplitude and computed the SSVEP signal based on the average of max electrode and its four surrounding electrodes. Moreover, we displayed the SSVEP amplitude for each foreground contrast without a background (black line) and with a background (red line) (Figure 3). As expected, the data demonstrate an increase of SSVEP amplitude with an increase of foreground contrast. We also demonstrated the surround suppression effect, which was measured as a relative reduction in amplitude of the SSVEP due to surround contrast. This is in line with a recent study, which originally developed and used this paradigm in healthy adults (Vanegas et al., 2015).

**Figure 3.**
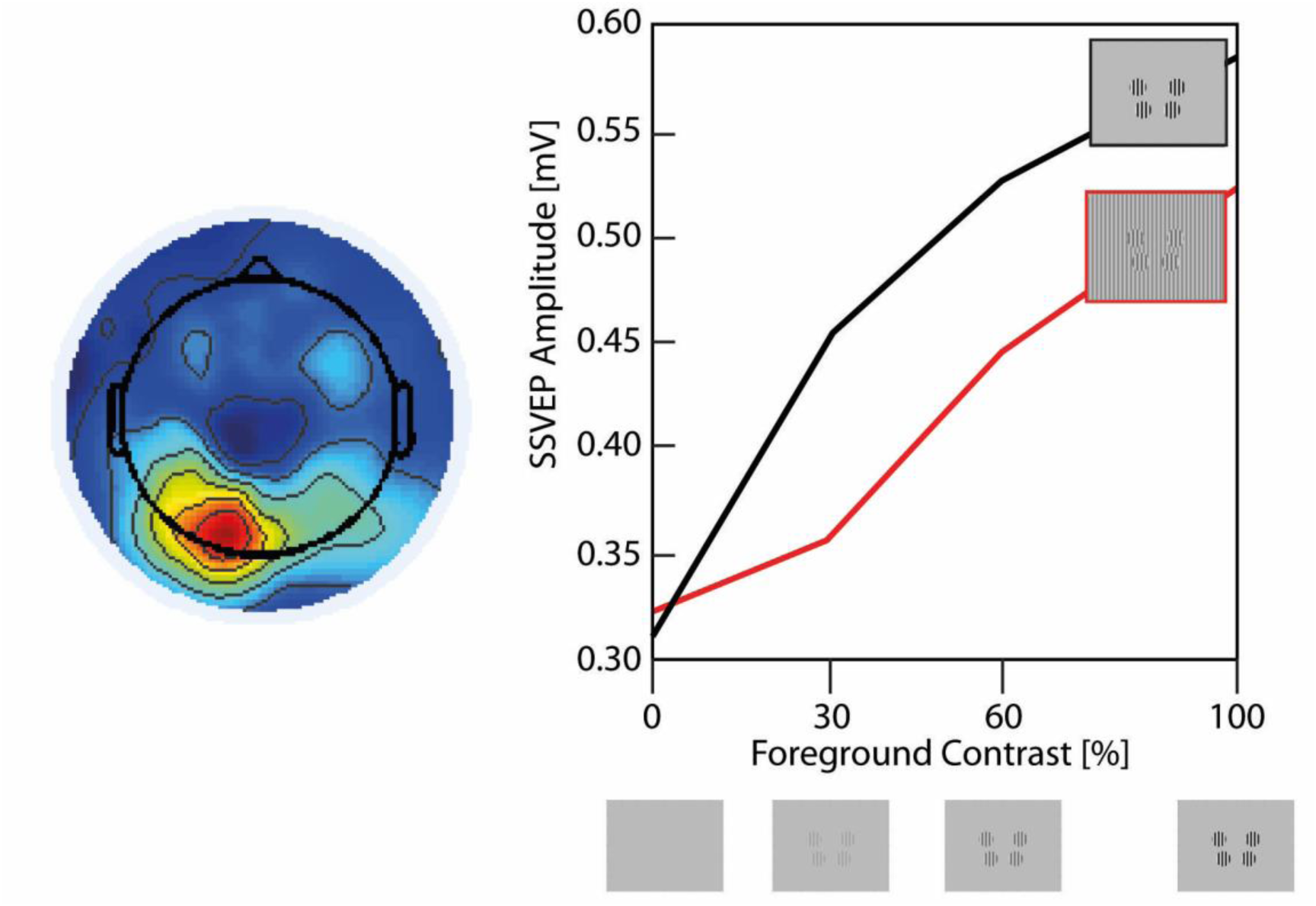
Surround Suppression Paradigm. The left plot displays the group average topographies of the 25 Hz steady-state visual evoked potential (SSVEP) amplitude for the mean of all foreground contrasts without a background. On the right panel, we displayed the SSVEP amplitude for each foreground contrast without a background (black line) and with background (red line).

### Naturalistic Stimuli Paradigm

Consistent with previously established methodologies, within-subject covariance matrices were computed across subjects and videos after standard EEG preprocessing. Thus, we obtained a set of component projections, which can be used as an ISC measure. We selected the three strongest correlated components and computed the corresponding correlation values separately for each component (Figure 4). It has been shown that the sum of the first three components explains sufficient variance of the data. The distribution of the ISC measure is in line with previous studies showing congruent distributions (Dmochowski et al., 2012; Dmochowski et al., 2014).

**Figure 4.**
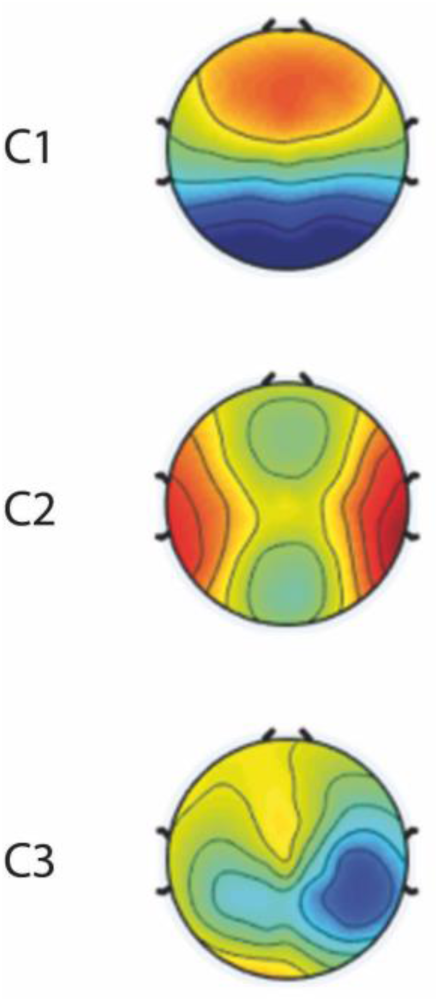
Naturalistic Stimuli Paradigm. Depicted are the scalp projections of the first three maximally correlated components (inter-subject correlation), averaged over all movies.

### Contrast Change Detection Paradigm

After standard EEG preprocessing, for each participant we merged the data from the three blocks. Target epochs were extracted from 500 ms before target onset to 1000 ms after peak sensory evidence. Moreover, response-locked epochs were extracted from 1000 ms before a response, to 300 ms after response button press. Trials were rejected if any scalp channel exceeded 100 μV. SSVEP (20 or 25 Hz depending on left or right target) based on stimulus-locked epochs and motor response signal (12.5–18 Hz) based on response-locked epochs were measured using the standard short-time Fourier transformation. The CPP analysis consisted simply of averaging the single-trial waveforms, which were baseline-corrected relative to the 500 ms interval before response onset. Figure 5 displays the sensory evidence encoding (SSVEP), the evidence accumulation (CPP), and motor preparation topographies. As expected, we found a posterior maximum for the SSVEP around the electrode Oz. The CPP signal shows the highest activity near the CPz electrode and the preparatory motor response signal (reduction in spectral amplitude relative to baseline) peaks over the electrodes C3 and C4.

**Figure 5.**
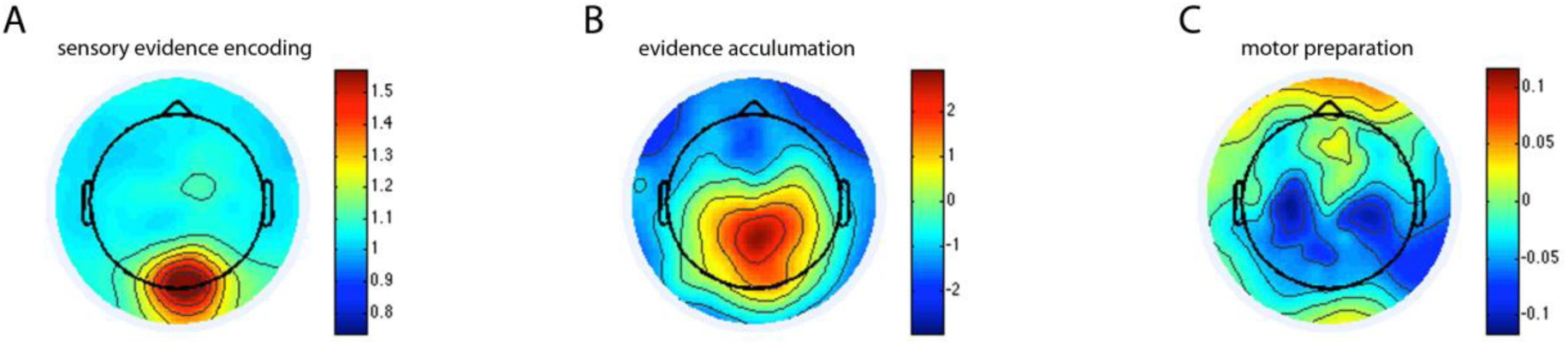
Contrast Change Detection Paradigm. The group average topographies are shown for the sensory evidence signal (represented as SSVEP, A), the response-locked CPP component reflecting evidence accumulation (B), and the decrease of beta frequency (12.5-18 Hz) spectral amplitude over left/right motor cortical areas at response relative to before target onset, reflecting motor preparation (C). Left-tilted and right-tilted targets are collapsed in all cases.

### Sequence Learning Paradigm

Following standard preprocessing, the EEG data were segmented based on stimulus onsets, which were each defined by the appearance of a filled white circle in one of the eight different spatial locations. Each epoch was 900 ms long (100 ms pre-stimulus to 800 post-stimulus presentation). For each of the five blocks, all artifact-free segments were extracted and subsequently baseline-corrected. For the technical validation, we averaged all trials within each block for each subject individually, and then calculated a group average. Although this averages across “still-unknown”, “newly-learned”, and “known” conditions within each block, the relative dominance of trial numbers in each category systematically varied over the course of the blocks as the sequence became better memorized. In Figure 6A, the ERP for the electrode CPz is depicted. We found a decrease in the P300 over the different blocks. In a next step, the behavioral “performance” was calculated as percentage correctly learned spatial locations, and the “learning rate” was defined as the newly learned locations in relation to all possible locations (Figure 6B). The corresponding P300 amplitude was calculated as the amplitude on electrode CPz with a latency of 350 – 500 ms post-stimulus presentation. We chose a slightly delayed time window, because recent studies have shown that the P300 in children exhibits a greater latency compared to adults (van Dinteren et al., 2014). Figure 6B reveals that while the performance is increasing with practice, the learning rate and the P300 are decreasing. These findings are in line with the assumption that the subjects learn to expect the order of the appearance of the targets over the course of the 5 blocks, which is represented by the performance and learning rate measure. In other words, over the course of the five blocks, the P300 amplitude decreases because the subjects are not as surprised by the appearance of the specific location of the stimulus presentation. Equivalent results have been reported in the original version of this paradigm from which the current one was adapted (Steinemann et al., 2016).

**Figure 6.**
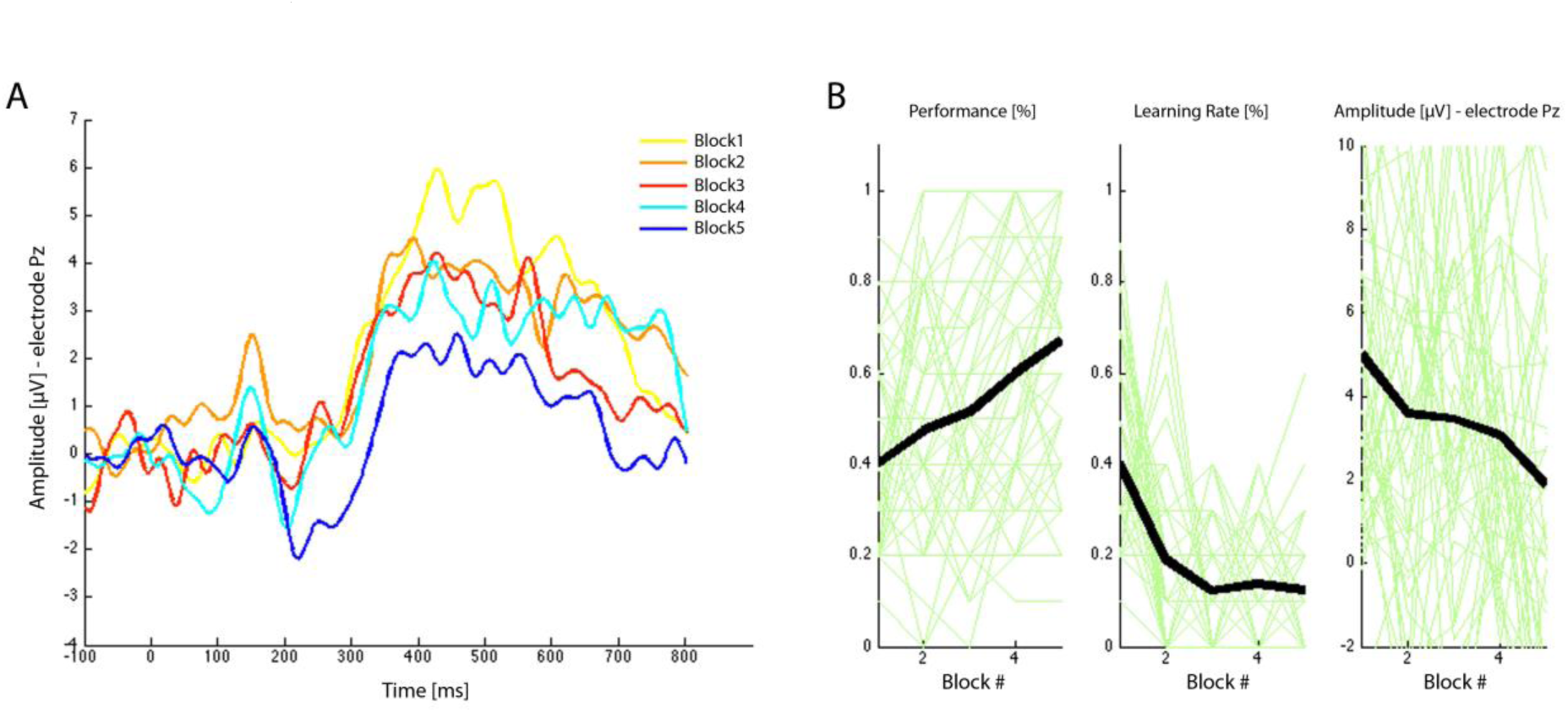
Sequence Learning Paradigm. In (A), the ERP for the electrode CPz is depicted for the average of each individual learning block. In (B), the P300 amplitude on electrode CPz, behavioral “performance” and “learning rate” are displayed. The black line indicates the mean of all subjects, whereas the green lines indicate each subject’s measures.

### Symbol Search Paradigm

For this paradigm, we mainly focused on the eye tracking data, but the newly established inventory of objective eye tracking measures can be complemented with topographic spatial and power analyses of the concurrently acquired EEG data. As described in the methods section, the goal of each trial is to determine whether either of two target symbols appears among a set of five search symbols. The presented graphic of the symbol search task was segmented into three subregions of interest: *targets, search group*, and *response buttons* (Figure 7A). From the eye tracking data, we calculated the number of saccade steps, number of repetitions, pupil size, and protracted gaze dwell times (fixation duration) for each subregion. Figure 7A displays all the fixations for a representative subject. The darkness of the color and the size of the circle indicate the duration of the fixations. Blue color indicates fixations outside of the current trial. Figure 7B represents the distribution of saccade amplitude, peak velocity and the angular histogram. In the second row of Figure 7B, the distribution of the durations of the fixations, the heat map and the allocation of the fixations are displayed. As expected, the data demonstrate that the eye tracker is able to track oculomotor activity while the subject is performing the task. This enables us to decompose the processing speed task into interpretable components of cognitive and perceptual processing, such as working memory, distractibility, uncertainty, and sustained attention.

**Figure 7.**
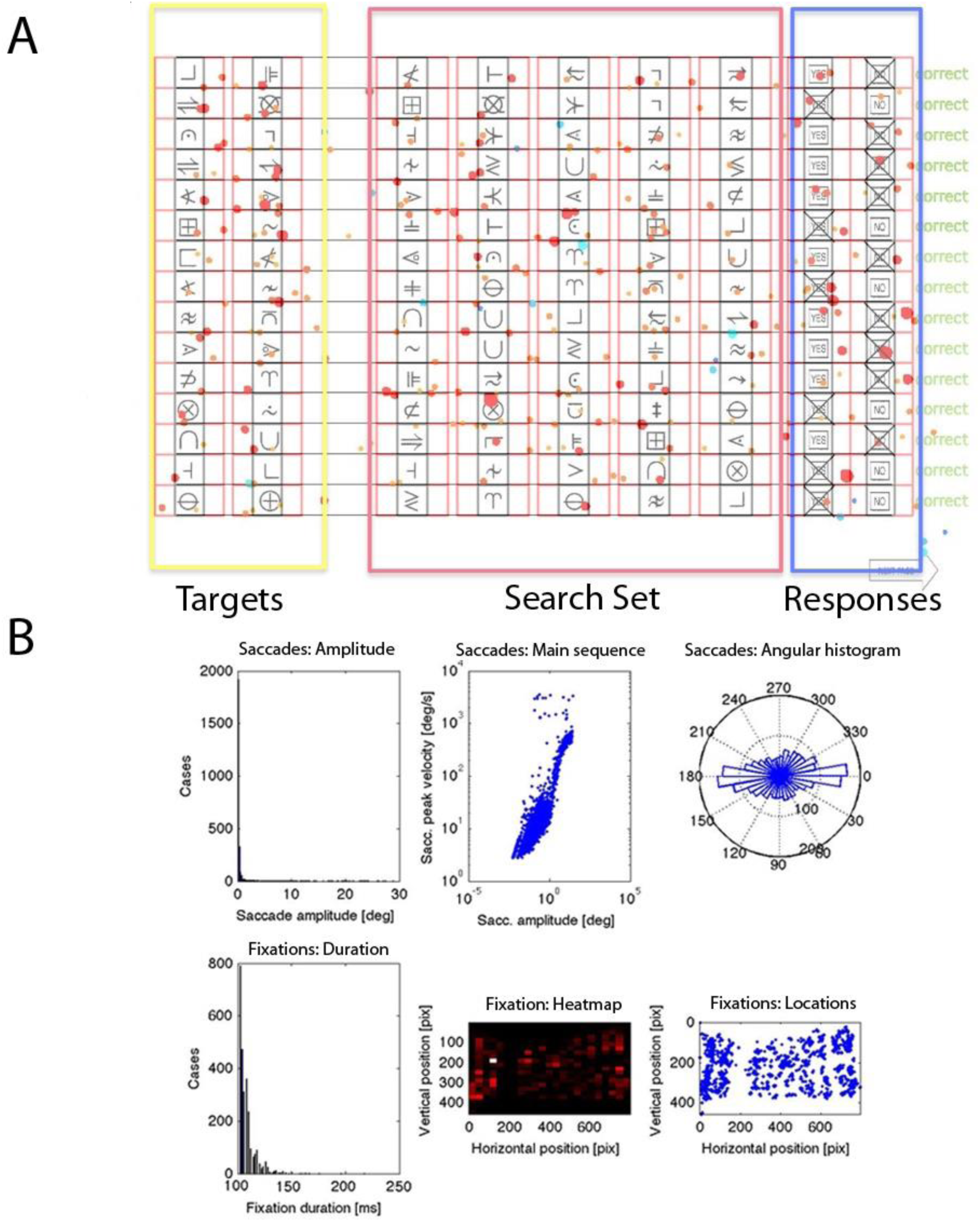
Symbol Search Paradigm. In (A), the three subregions of interest, *targets, search set* and *response buttons*, are displayed with all fixations for a representative subject superimposed. The darkness of the color and the size of the circle indicate the duration of the fixations. Blue color indicates fixations outside of the current trial. (B) represents the distribution of saccade amplitude, peak velocity and the angular histogram. In the second row, the distribution of the durations of the fixations, the heat map and the allocation of the fixations are displayed.

## Acknowledgem Ents

The data resource and work presented here were supported by gifts to the Child Mind Institute from Phyllis Green, Randolph Cowen, and Joseph Healey. Nicolas Langer was supported through the CMI Endeavor Scientists Program. Michael Milham is a Randolph Cowen and Phyllis Green Scholar.

We would like to thank the participants and their families for taking the time to be in the study, as well as their willingness to have their data shared with the scientific community. We would also like to thank the members of the Mind Research Network, particularly Margaret King and Vince Calhoun, for their support in using the COINS neuroinformatics platform.

## Author Contributions

NL: Experiment Design, Data Organization, Data Analysis, Drafting of Manuscript

EH: Data Acquisition, Data Organization, Data Analysis, Drafting of Manuscript

LA: Data Acquisition, Data Organization, Data Analysis, Drafting of Manuscript

HX: Data Acquisition, Data Organization, Data Analysis, Drafting of Manuscript

RJ: Experiment Design, Data Organization, Data Analysis, Drafting of Manuscript

SH: Experiment Design and Data Analysis SC: Experiment Design, Drafting of Manuscript

EM: Data Acquisition

LP: Experiment Design, Drafting of Manuscript

MM: Experiment Design, Drafting of Manuscript

SP: Experiment Design, Drafting of Manuscript

## Competing Interests

None of the authors have a conflict of interest to report.

## References

Annett M (1970) A classification of hand preference by association analysis. Br J Psychol 61:303–321.

Arns M, Conners CK, Kraemer HC (2013) A decade of EEG Theta/Beta Ratio Research in ADHD: a meta analysis. Journal of attention disorders 17:374–383.

Barry RJ, Clarke AR, Johnstone SJ, Brown CR (2009) EEG differences in children between eyes-closed and eyes-open resting conditions. Clinical neurophysiology: official journal of the International Federation of Clinical Neurophysiology 120:1806–1811.

Bartels A, Zeki S (2004) Functional brain mapping during free viewing of natural scenes. Human brain mapping 21:75–85.

Battista J, Badcock DR, McKendrick AM (2011) Migraine increases centre-surround suppression for drifting visual stimuli. PloS one 6:e18211.

Ben-Yakov A, Honey CJ, Lerner Y, Hasson U (2012) Loss of reliable temporal structure in event-related averaging of naturalistic stimuli. NeuroImage 63:501–506.

Biswal B, Yetkin FZ, Haughton VM, Hyde JS (1995) Functional connectivity in the motor cortex of resting human brain using echo-planar MRI. Magnetic resonance in medicine 34:537–541.

Borbely AA, Achermann P (1999) Sleep homeostasis and models of sleep regulation. Journal of biological rhythms 14:557–568.

Brainard DH (1997) The Psychophysics Toolbox. Spatial vision 10:433–436.

Burgess A, Flint J, Adshead H (1992) Factor structure of the Wechsler Adult Intelligence Scale-revised (WAIS-R): a clinical sample. The British journal of clinical psychology / the British Psychological Society 31 (Pt 3):336–338.

Cantlon JF, Li R (2013) Neural activity during natural viewing of Sesame Street statistically predicts test scores in early childhood. PLoS biology 11:e1001462.

Cavanaugh JR, Bair W, Movshon JA (2002) Nature and Interaction of Signals From the Receptive Field Center and Surround in Macaque V1 Neurons. Journal of neurophysiology 88:2530–2546.

Chen AC, Feng W, Zhao H, Yin Y, Wang P (2008) EEG default mode network in the human brain: spectral regional field powers. NeuroImage 41:561–574.

Cohen S, Parra LC (2016) Memorable audiovisual narratives synchronize sensory and supramodal neural responses. eNeuro 3.

Cuthbert BN, Insel TR (2013) Toward the future of psychiatric diagnosis: the seven pillars of RDoC. BMC medicine 11:126.

Dakin S, Carlin P, Hemsley D (2005) Weak suppression of visual context in chronic schizophrenia. Current biology: CB 15:R822–824.

Damoiseaux JS, Rombouts SA, Barkhof F, Scheltens P, Stam CJ, Smith SM, Beckmann CF (2006) Consistent resting-state networks across healthy subjects. Proceedings of the National Academy of Sciences of the United States of America 103:13848–13853.

De Vico Fallani F, Maglione A, Babiloni F, Mattia D, Astolfi L, Vecchiato G, De Rinaldis A, Salinari S, Pachou E, Micheloyannis S (2010) Cortical network analysis in patients affected by schizophrenia. Brain topography 23:214–220.

Deuker L, Bullmore ET, Smith M, Christensen S, Nathan PJ, Rockstroh B, Bassett DS (2009) Reproducibility of graph metrics of human brain functional networks. NeuroImage 47:14601468.

Dmochowski JP, Sajda P, Dias J, Parra LC (2012) Correlated components of ongoing EEG point to emotionally laden attention – a possible marker of engagement? Frontiers in human neuroscience 6:112.

Dmochowski JP, Bezdek MA, Abelson BP, Johnson JS, Schumacher EH, Parra LC (2014) Audience preferences are predicted by temporal reliability of neural processing. Nature communications 5:4567.

Dockree PM, Kelly SP, Robertson IH, Reilly RB, Foxe JJ (2005) Neurophysiological markers of alert responding during goal-directed behavior: a high-density electrical mapping study. NeuroImage 27:587–601.

Dockree PM, Kelly SP, Foxe JJ, Reilly RB, Robertson IH (2007) Optimal sustained attention is linked to the spectral content of background EEG activity: greater ongoing tonic alpha (approximately 10 Hz) power supports successful phasic goal activation. The European journal of neuroscience 25:900907.

Donchin E (1981) Presidential address, 1980. SurpriseL.Surprise? Psychophysiology 18:493–513.

Donders J, Tulsky DS, Zhu J (2001) Criterion validity of new WAIS-II subtest scores after traumatic brain injury. Journal of the International Neuropsychological Society: JINS 7:892–898.

Donner TH, Siegel M, Fries P, Engel AK (2009) Buildup of choice-predictive activity in human motor cortex during perceptual decision making. Current biology: CB 19:1581–1585.

Duering M, Gesierich B, Seiler S, Pirpamer L, Gonik M, Hofer E, Jouvent E, Duchesnay E, Chabriat H, Ropele S, Schmidt R, Dichgans M (2014) Strategic white matter tracts for processing speed deficits in age-related small vessel disease. Neurology 82:1946–1950.

Eckert MA (2011) Slowing down: age-related neurobiological predictors of processing speed. Frontiers in neuroscience 5:25.

Fernandez T, Harmony T, Rodriguez M, Bernal J, Silva J, Reyes A, Marosi E (1995) EEG activation patterns during the performance of tasks involving different components of mental calculation. Electroencephalography and clinical neurophysiology 94:175–182.

Fontanini A, Katz DB (2008) Behavioral states, network states, and sensory response variability. Journal of neurophysiology 100:1160–1168.

Foss-Feig JH, Tadin D, Schauder KB, Cascio CJ (2013) A substantial and unexpected enhancement of motion perception in autism. The Journal of neuroscience: the official journal of the Society for Neuroscience 33:8243–8249.

Fox MD, Greicius M (2010) Clinical applications of resting state functional connectivity. Frontiers in systems neuroscience 4:19.

Friston KJ, Frith CD, Liddle PF, Frackowiak RS (1993) Functional connectivity: the principal-component analysis of large (PET) data sets. Journal of cerebral blood flow and metabolism: official journal of the International Society of Cerebral Blood Flow and Metabolism 13:5–14.

Gevins A, Smith ME, McEvoy L, Yu D (1997) High-resolution EEG mapping of cortical activation related to working memory: effects of task difficulty, type of processing, and practice. Cerebral cortex 7:374–385.

Golomb JD, McDavitt JR, Ruf BM, Chen JI, Saricicek A, Maloney KH, Hu J, Chun MM, Bhagwagar Z (2009) Enhanced visual motion perception in major depressive disorder. The Journal of neuroscience: the official journal of the Society for Neuroscience 29:9072–9077.

Gratton G, Coles MG, Sirevaag EJ, Eriksen CW, Donchin E (1988) Pre- and poststimulus activation of response channels: a psychophysiological analysis. Journal of experimental psychology Human perception and performance 14:331–344.

Hanson SJ, Gagliardi AD, Hanson C (2009) Solving the brain synchrony eigenvalue problem: conservation of temporal dynamics (fMRI) over subjects doing the same task. Journal of computational neuroscience 27:103–114.

Hasson U, Malach R, Heeger DJ (2010) Reliability of cortical activity during natural stimulation. Trends in cognitive sciences 14:40–48.

Hasson U, Nir Y, Levy I, Fuhrmann G, Malach R (2004) Intersubject synchronization of cortical activity during natural vision. Science 303:1634–1640.

Hasson U, Furman O, Clark D, Dudai Y, Davachi L (2008) Enhanced intersubject correlations during movie viewing correlate with successful episodic encoding. Neuron 57:452–462.

Insel TR, Cuthbert BN (2015) Medicine. Brain disorders? Precisely. Science 348:499–500.

Ivonin AA, Tsitseroshin MN, Pogosyan AA, Shuvaev VT (2004) Genetic determination of neurophysiological mechanisms of cortical-subcortical integration of bioelectrical brain activity. Neuroscience and behavioral physiology 34:369–378.

Jeste SS, Frohlich J, Loo SK (2015) Electrophysiological biomarkers of diagnosis and outcome in neurodevelopmental disorders. Current opinion in neurology 28:110–116.

John ER, Prichep LS, Fridman J, Easton P (1988) Neurometrics: computer-assisted differential diagnosis of brain dysfunctions. Science 239:162–169.

John ER, Ahn H, Prichep L, Trepetin M, Brown D, Kaye H (1980) Developmental equations for the electroencephalogram. Science 210:1255–1258.

Joy S, Kaplan E, Fein D (2004) Speed and memory in the WAIS-III Digit Symbol-Coding subtest across the adult lifespan. Archives of clinical neuropsychology: the official journal of the National Academy of Neuropsychologists 19:759–767.

Kapur S, Phillips AG, Insel TR (2012) Why has it taken so long for biological psychiatry to develop clinical tests and what to do about it? Molecular psychiatry 17:1174–1179.

Karis D, Druckman D, Lissak R, Donchin E (1984) A psychophysiological analysis of bargaining. ERPs and facial expressions. Annals of the New York Academy of Sciences 425:230–235.

Kitsune GL, Cheung CH, Brandeis D, Banaschewski T, Asherson P, McLoughlin G, Kuntsi J (2015) A Matter of Time: The Influence of Recording Context on EEG Spectral Power in Adolescents and Young Adults with ADHD. Brain topography 28:580–590.

Kondacs A, Szabo M (1999) Long-term intra-individual variability of the background EEG in normals. Clinical neurophysiology: official journal of the International Federation of Clinical Neurophysiology 110:1708–1716.

Kozak MJ, Cuthbert BN (2016) The NIMH Research Domain Criteria Initiative: Background, Issues, and Pragmatics. Psychophysiology 53:286–297.

Langer N, Pedroni A, Gianotti LR, Hanggi J, Knoch D, Jancke L (2012) Functional brain network efficiency predicts intelligence. Human brain mapping 33:1393–1406.

Lauritzen TZ, Ales JM, Wade AR (2010) The effects of visuospatial attention measured across visual cortex using source-imaged, steady-state EEG. Journal of vision 10.

Lehmann D, Ozaki H, Pal I (1987) EEG alpha map series: brain micro-states by space-oriented adaptive segmentation. Electroencephalography and clinical neurophysiology 67:271–288.

Lehmann D, Strik WK, Henggeler B, Koenig T, Koukkou M (1998) Brain electric microstates and momentary conscious mind states as building blocks of spontaneous thinking: I. Visual imagery and abstract thoughts. International journal of psychophysiology: official journal of the International Organization of Psychophysiology 29:1–11.

Levitt JB, Lund JS (1997) Contrast dependence of contextual effects in primate visual cortex. Nature 387:73–76.

Lezak MD (1995) Neuropsychological assessment. New York: Oxford University Press.

Lin Z, Chen M, Ma Y (2010) The Augmented Lagrange Multiplier Method for Exact Recovery of Corrupted Low-Rank Matrices. arXiv:10095055.

Linkenkaer-Hansen K, Smit DJ, Barkil A, van Beijsterveldt TE, Brussaard AB, Boomsma DI, van Ooyen A, de Geus EJ (2007) Genetic contributions to long-range temporal correlations in ongoing oscillations. The Journal of neuroscience: the official journal of the Society for Neuroscience 27:13882–13889.

Loo SK, Lenartowicz A, Makeig S (2015) Research Review: use of EEG biomarkers in child psychiatry research – current state and future directions. Journal of child psychology and psychiatry, and allied disciplines.

Macpherson H, Pipingas A, Silberstein R (2009) A steady state visually evoked potential investigation of memory and ageing. Brain and cognition 69:571–579.

Mars RB, Debener S, Gladwin TE, Harrison LM, Haggard P, Rothwell JC, Bestmann S (2008) Trial-by-trial fluctuations in the event-related electroencephalogram reflect dynamic changes in the degree of surprise. The Journal of neuroscience: the official journal of the Society for Neuroscience 28:12539–12545.

Michel C, Koenig T, Brandeis D, Gianotti LR, Wackermann J (2009) Electrical Neuroimaging. Cambridge: Cambridge University Press.

Moisello C, Meziane HB, Kelly S, Perfetti B, Kvint S, Voutsinas N, Blanco D, Quartarone A, Tononi G, Ghilardi MF (2013) Neural activations during visual sequence learning leave a trace in post-training spontaneous EEG. PloS one 8:e65882.]

Murray MM, Brunet D, Michel CM (2008) Topographic ERP analyses: a step-by-step tutorial review. Brain topography 20:249–264.

Napflin M, Wildi M, Sarnthein J (2007) Test-retest reliability of resting EEG spectra validates a statistical signature of persons. Clinical neurophysiology: official journal of the International Federation of Clinical Neurophysiology 118:2519–2524.

Neville HJ, Kutas M, Chesney G, Schmidt AL (1996) Event-related brain potentials during initial encoding and recognition memory of congruous and incongruous words. Journal of Memory and Language 25:75–92.

O’Connell RG, Dockree PM, Kelly SP (2012) A supramodal accumulation-to-bound signal that determines perceptual decisions in humans. Nature neuroscience 15:1729–1735.

Orekhova EV, Stroganova TA, Posikera IN, Malykh SB (2003) Heritability and “environmentability” of electroencephalogram in infants: the twin study. Psychophysiology 40:727–741.

Paller KA, Kutas M, Shimamura AP, Squire LR (1987) Brain responses to concrete and abstract words reflect processes that correlate with later performance on a test of stem-completion priming. Electroencephalography and clinical neurophysiology Supplement 40:360–365.

Pascual-Marqui R (2007) Instanteneous and lagged measurements of linear and nonlinear dependence between groups of multivariate times series: frequency decomposition. arXiv:07111455[statME].

Pascual-Marqui RD, Michel CM, Lehmann D (1995) Segmentation of brain electrical activity into microstates: model estimation and validation. IEEE transactions on bio-medical engineering 42:658–665.

Pelli DG (1997) The VideoToolbox software for visual psychophysics: transforming numbers into movies. Spatial vision 10:437–442.

Perrin F, Pernier J, Bertrand O, Giard MH, Echallier JF (1987) Mapping of Scalp Potentials by Surface Spline Interpolation. Electroencephalography and clinical neurophysiology 66:75–81.

Petroni A, Cohen S, Langer N, Henin S, Vanderwal T, Milham MP, Parra LC (2016) Age and sex affect intersubject correlation of EEG throughout development. bioArxiv 089060.

Pfurtscheller G, Neuper C, Ramoser H, Muller-Gerking J (1999) Visually guided motor imagery activates sensorimotor areas in humans. Neuroscience letters 269:153–156.

Posthuma D, Neale MC, Boomsma DI, de Geus EJ (2001) Are smarter brains running faster? Heritability of alpha peak frequency, IQ, and their interrelation. Behavior genetics 31:567–579.

Regan D (1966) An effect of stimulus colour on average steady-state potentials evoked in man. Nature 210:1056–1057.

Regan D (1989) Human Brain Electrophysiology: Evoked Potentials and Evoked Magnetic Fields in Science and Medicine. New York: Elsevier.

Royer FL, Gilmore GC, Gruhn JJ (1981) Normative data for the Symbol Digit Substitution Task. Journal of clinical psychology 37:608–614.

Salthouse TA, Ferrer-Caja E (2003) What needs to be explained to account for age-related effects on multiple cognitive variables? Psychology and aging 18:91–110.

Schacter DL, Wagner AD (1999) Perspectives: neuroscience. Remembrance of things past. Science 285:1503–1504.

Seymour K, Stein T, Sanders LL, Guggenmos M, Theophil I, Sterzer P (2013) Altered contextual modulation of primary visual cortex responses in schizophrenia. Neuropsychopharmacology: official publication of the American College of Neuropsychopharmacology 38:2607–2612.

Smallwood J, Nind L, O’Connor RC (2009) When is your head at? An exploration of the factors associated with the temporal focus of the wandering mind. Consciousness and cognition 18:118–125.

Smallwood J, McSpadden M, Luus B, Schooler J (2008) Segmenting the stream of consciousness: the psychological correlates of temporal structures in the time series data of a continuous performance task. Brain and cognition 66:50–56.

Smith SM, Fox PT, Miller KL, Glahn DC, Fox PM, Mackay CE, Filippini N, Watkins KE, Toro R, Laird AR, Beckmann CF (2009) Correspondence of the brain’s functional architecture during activation and rest. Proceedings of the National Academy of Sciences of the United States of America 106:13040–13045.

Steinemann NA, Moisello C, Ghilardi MF, Kelly SP (2016) Tracking neural correlates of successful learning over repeated sequence observations. NeuroImage 137:152–164.

Stephens GJ, Silbert LJ, Hasson U (2010) Speaker-listener neural coupling underlies successful communication. Proceedings of the National Academy of Sciences of the United States of America 107:14425–14430.

Twomey DM, Murphy PR, Kelly SP, O’Connell RG (2015) The classic P300 encodes a build-to-threshold decision variable. The European journal of neuroscience 42:1636–1643.

van Dinteren R, Arns M, Jongsma ML, Kessels RP (2014) P300 development across the lifespan: a systematic review and meta-analysis. PloS one 9:e87347.

Van Rooy C, Stough C, Pipingas A, Hocking C, Silberstein RB (2001) Spatial working memory and intelligence biological correlate. Intelligence 29:275–292.

Vanderwal T, Kelly C, Eilbott J, Mayes LC, Castellanos FX (2015) Inscapes: A movie paradigm to improve compliance in functional magnetic resonance imaging. NeuroImage 122:222–232.

Vanegas MI, Blangero A, Kelly SP (2013) Exploiting individual primary visual cortex geometry to boost steady state visual evoked potentials. Journal of neural engineering 10:036003.

Vanegas MI, Blangero A, Kelly SP (2015) Electrophysiological indices of surround suppression in humans. Journal of neurophysiology 113:1100–1109.

Vialatte FB, Maurice M, Dauwels J, Cichocki A (2010) Steady-state visually evoked potentials: focus on essential paradigms and future perspectives. Progress in neurobiology 90:418–438.

Vogel F (2000) Genetics and the Electroencephalogram. Berlin: Springer.

Wagner AD, Koutstaal W, Schacter DL (1999) When encoding yields remembering: insights from event-related neuroimaging. Philosophical transactions of the Royal Society of London Series B, Biological sciences 354:1307–1324.

Wechsler D (2004) The Wechsler intelligence scale for children, fourth edition Edition. London: Pearson.

Xing J, Heeger DJ (2000) Center-surround interactions in foveal and peripheral vision. Vision research 40:3065–3072.

Zacks JM, Tversky B (2001) Event structure in perception and conception. Psychological bulletin 127:321.

Zenger-Landolt B, Heeger DJ (2003) Response suppression in v1 agrees with psychophysics of surround masking. The Journal of neuroscience: the official journal of the Society for Neuroscience 23:6884–6893.

